# Disease as a Mediator of Somatic Mutation - Lifespan Coevolution: Model Analysis and Empirical Tests

**DOI:** 10.1101/2025.02.05.636604

**Authors:** Michael E. Hochberg

## Abstract

Multicellular organisms are confronted not only with germline mutations, but also with mutations emerging in somatic cells. Somatic mutations can lead to diverse pathologies, including cancer. Somatic mutation rate (SMR) can be constrained by evolved protection mechanisms, notably those repairing damaged DNA, or eliminating or slowing the growth of mutated cells. However, in a broader context, life history traits such as body mass, age of first reproduction and reproductive lifespan can also be subject to selection due to the negative fitness impacts of disease. Here, I analyze a simple coevolutionary model of somatic mutation rate and lifespan. The trait dynamics are formulated as an evolutionary-response system through an explicit, frequency-dependent invasion-fitness function. Evolution in the model is driven in part by the negative fitness impacts of disease, represented phenomenologically as increasing with both SMR and lifespan. I investigate relations between selective forces and (co)evolutionary responses, notably showing the possibility of either monotone or oscillatory non-equilibrium dynamics and fast or slow returns to equilibrium. Model predictions are then compared to recently published data on body mass, lifespan and somatic mutation rate. I show that the model (*1*) can explain the non-linear empirical relationship between somatic mutation and lifespan, (*2*) predicts the evolution of longer lifespans through a heretofore ignored feedback loop, (*3*) predicts opposite body-mass scaling of cumulative mutational burden and lifespan-normalized mutation rate, and (*4*) is compatible with observed linear mutation accumulation with age reflecting a hypothesized washing-out of highly mutated cells. I discuss the findings and their generality to somatic diseases, notably cancers and to aging itself.

## Introduction

Somatic mutation is observed throughout multicellular organisms, including animals, plants and fungi. Mutations can occur from the first cell divisions, through juvenile development and growth, and during adulthood in non-dividing cells or during cell divisions that replace aged, damaged or dead cells. Much of our current understanding of somatic mutation comes from studies of its cumulative burden in humans (Lynch, 2010b), its causes and impacts, including genome maintenance (MacRae et al., 2015; Mustjoki and Young, 2021; Zhang et al., 2021; Vijg, 2021), aging (Vijg, 2021; Schumacher et al., 2021; Chatsirisupachai and De Magalhães, 2024), the immune system (Marongiu and DeGregori, 2022), and the origin and progression of conditions and pathologies, including somatic mosaicism (Mustjoki and Young, 2021; Hall, 1988; Frank, 2010, 2014; D’Gama and Walsh, 2018; Vijg and Dong, 2020) and cancer (Frank, 2010, 2005; Laconi et al., 2020). Relatedly, studies of the ecological and evolutionary dynamics of somatic mutation have gained attention with research focusing on aging (Laconi et al., 2020; Tian et al., 2017), clonal expansions (Mustjoki and Young, 2021; Chatsirisupachai and De Magalhães, 2024; Marongiu and DeGregori, 2022; Forsberg et al., 2017; Olafsson and Anderson, 2021; Evans and DeGregori, 2021; Herms and Jones, 2023), disease (Frank, 2014; Laconi et al., 2020), and life history traits (Kokko and Hochberg, 2015; Cagan et al., 2022). An outstanding question emerging from these and other studies is the extent to which selection acts on mutation-derived disease risk, either directly through genomic and cellular maintenance or downstream disease prevention, or indirectly through life history evolution.

Somatic mutations can manifest in several non-mutually exclusive ways. They may occur in nuclear or mitochondrial DNA (Kopinski et al., 2021; Harrington et al., 2023), emerge in otherwise healthy cell lineages either during organism development (Frank, 2010; D’Gama and Walsh, 2018; De, 2011; Rockweiler et al., 2023) or tissue renewal or regeneration (Zhu et al., 2019; Martin et al., 2024), be limited to a single cell or small group of cells (Lee-Six et al., 2019), or they may expand and accumulate during, or contribute to, oncogenesis and tumorigenesis (Evans and DeGregori, 2021; Herms and Jones, 2023; Wu et al., 2018; Rozhok and DeGregori, 2020; Weeden et al., 2023). Although exceptions exist (discussed in Marongiu and DeGregori, 2022; Colom et al., 2021), single-cell or small-lineage mutations that remain confined, whether because of protective mechanisms (e.g., DNA repair, apoptosis, innate immune surveillance, cell turnover and cell-cell competition (Cairns, 1975; Bowling et al., 2019), or tissue architecture (Herms and Jones, 2023) or because they fail to expand, are expected to have limited impacts on tissue structure and organ system function (Evans and DeGregori, 2021; Hall et al., 2019). In contrast to confined mutational effects, intrinsic (e.g., inflammation, aging) or extrinsic (e.g., UV radiation, lifestyle-associated exposures) conditions can favor the expansion of mutated cells and increase risks of disease-associated morbidity and mortality (Weeden et al., 2023; Hochberg and Noble, 2017).

Mutations conferring cellular gain-of-function phenotypes or promoting clonal expansions (Evans and DeGregori, 2021; Hall et al., 2019; Martincorena, 2019; Kakiuchi and Ogawa, 2021), and specifically those contributing to invasive carcinoma, illustrate how mutations can produce pathologies years or decades after the first genetic lesions in humans (Gerstung et al., 2020). Here, genomic and cellular dynamics are influenced by host environments, including the pathology itself (Laconi et al., 2020), the surrounding aging microenvironment and tissues (Fane and Weeraratna, 2020), and more indirectly, lifestyle-associated exposures (Wu et al., 2018; Weeden et al., 2023). Physiological aging, used here as progressive decline in function, can also mediate the downstream effects of somatic mutation, though the causal effects of mutations on aging phenotypes are controversial (Vijg, 2021; Schumacher et al., 2021; Chatsirisupachai and De Magalhães, 2024; Franco et al., 2022; Shen et al., 2024). Evidence suggests that, after birth, somatic mutations in many human tissues accumulate approximately linearly with chronological age, implying relatively stable tissue-specific accumulation rates rather than strong age-dependent acceleration (Manders et al., 2021; Alexandrov et al., 2015). Aging nevertheless has been hypothesized to exacerbate the clinical manifestations of certain genetic or epigenetic cellular changes. These changes may increase disease risk (Shen et al., 2024; Niccoli and Partridge, 2012; Franceschi et al., 2018) and are associated with increases in the population incidence and individual severity of certain diseases, including cancers (Hochberg and Noble, 2017; St Sauver et al., 2015; DeGregori et al., 2020).

Evolutionary theories suggest that the accumulation of deleterious cell variants, cell lineages (e.g., somatic mosaicism) and cell aggregates (e.g., neoplasms) will be influenced by age-specific selection on their negative fitness effects (Medawar, 1952; Williams, 1957; Rozhok and DeGregori, 2016). The extent to which selection acts on genomic and cellular maintenance and/or mechanisms that attenuate downstream mutation-associated disease is little explored, but somatic mutation rates should be shaped by natural selection primarily when their associated fitness effects occur at ages that influence survival or reproduction. For example, in humans, some estimates of prenatal somatic mutation rates are elevated (Manders et al., 2021; Kuijk et al., 2019), whereas after birth mutations appear to accumulate approximately linearly at tissue-specific rates (Manders et al., 2021). These observations do not by themselves demonstrate relaxed organism-level selection; rather, they indicate ontogenetic differences in mutational processes and genome surveillance that can interact with age-specific selection (Stearns, 1987; Krakauer and Mira, 1999; Baudisch, 2005). Other studies have considered how various factors could lead to age-dependence in disease resistance and tolerance (e.g., Hochberg et al., 2013; Nunney, 2013; Buckingham and Ashby, 2024). A theoretical expectation is that selection on heritable variants whose effects are expressed before or during reproduction is generally stronger than selection on effects expressed late in life, although the precise age profile depends on demography and trait effects (Cheng and Kirkpatrick, 2021). In this perspective, pleiotropic effects and time lags between somatic mutations and their downstream effects can explain why mutations may persist to some extent at young ages, and why associated conditions and diseases may be weakly opposed by selection at sufficiently advanced ages (Rozhok and DeGregori, 2020; Gladyshev, 2016).

Theories to explain the evolution of somatic mutation rate (SMR) generally invoke negative fitness effects of mutations (Lynch, 2010a) and/or costs of DNA repair (Kirkwood, 2005). Evolution of germline-encoded mechanisms that lower SMR is expected to occur when negative effects are manifested at ages where the force of selection is sufficient to offset costs of enhanced protection (Kirkwood and Holliday, 1979; Campisi, 2005). Age-related cancer incidence, for example, often increases as the force of selection declines (Rozhok and DeGregori, 2020; Hochberg and Noble, 2017), consistent with selection reducing the incidence or severity of fitness-threatening cancers at ages of high reproductive value. This is despite the frequent presence of cancer-associated mutations in otherwise normal cells (e.g., Martincorena and Campbell, 2015) and some tumors that have little effect on survival and reproduction. Moreover, given the large number of cell divisions during a lifetime in vertebrates such as mammals, birds, and some reptiles, together with the expectation that many mutations are either of negligible effect or are deleterious and occur in short-lived cells, a prediction is that costly genomic maintenance will be imperfect, and will be supplemented by mechanisms focused on the more direct costs to survival and reproduction due to negative downstream effects (see also Frank, 2023).

Evolutionary responses to somatic disruption can extend beyond proximal mechanisms such as DNA repair, telomere maintenance, and apoptosis (Tian et al., 2018; Risques and Promislow, 2018), to life history traits such as body size (Rozhok and DeGregori, 2019; Erten et al., 2020; Avila and Lehmann, 2023) and reproduction (Kokko and Hochberg, 2015; Hochberg et al., 1992; Kapsetaki et al., 2024). Support for associations between mutation and life history variables comes from several sources. Consistent with early somatic-mutation theories of aging and carcinogenesis (Failla, 1958), comparative data show an inverse relationship between SMR and lifespan across species (Cagan et al., 2022), with metabolic rates as a possible mediator (Dang, 2015). Similarly, there is evidence for a positive relationship between DNA repair and body size across species (Promislow, 1994). In contrast, evidence for associations between DNA repair and lifespan is supported for some within-species comparisons (Freitas and De Magalhães, 2011), but remains equivocal across species (Promislow, 1994). Despite these empirical patterns, limited attention has been given to testable theory on the coevolution of mutation rate and life history traits (but see Rozhok and DeGregori, 2019; Avila and Lehmann, 2023).

Here, I develop theory for how pathologies stemming from somatic mutation -- and cancer in particular -- can shape not only SMRs but also life history in the form of lifespan, defined as the typical age at which the force of selection becomes insignificant. My objective is to elucidate the key processes driving three central concepts in organismal biology: mutation rate, life history, and disease. We make no claim that disease is the primary determinant of somatic mutation rate. The model deliberately isolates the disease-mediated component of somatic-mutation-rate and life-history coevolution, ignoring other determinants (in particular germline mutation rate and the shared replication- and repair-fidelity machinery with which somatic mutation rate co-varies) whose evolution may account for some of the observed cross-species variation in somatic mutation rate (see Discussion). The present study therefore takes an initial step in developing and analyzing a simple phenomenological mathematical model of SMR-lifespan coevolution. I focus on equilibrium and non-equilibrium model conditions to make predictions of how different selection scenarios could translate into single-species dynamics and cross-species phylogenetic patterns. I then compare model predictions with data from the comparative study by Cagan et al.(2022), the results of which provide insights into possible links between mutation rate, life history and cancer. I conclude by proposing a hypothesized mechanism by which cancer could contribute to the evolution of lifespan in multicellular organisms. I also discuss how the framework may generalize to the coevolution between homeostatic mechanisms, aging, and life history.

### Model Assumptions

We develop a simple differential equation model as a first step in analyzing phenotypic evolution of SMR and lifespan. The model is phenomenological, that is, relations between trait evolution and selective drivers are assumed to take the form of constants or linear or bilinear functions, motivated by empirical considerations. We model downstream costs and buffering of somatic mutation with bilinear terms because the fitness consequences of somatic variants are expected to depend both on mutational burden and on the time available for clonal persistence, expansion, and pathology. These terms are phenomenological first-order approximations rather than empirically established functional forms. Thus, the patterns and processes generated by the present model are preliminary, and will need to be tested in future studies.

Our core assumption that SMR and lifespan are evolutionarily linked is motivated by data on age-related (i) accumulation of somatic mutations (Manders et al., 2021; Martincorena and Campbell, 2015; Blokzijl et al., 2016; Moore et al., 2021), (ii) increases in mutation-associated diseases including cancers (Marongiu and DeGregori, 2022; Proukakis, 2020; Evans and Walsh, 2023), (iii) decreases in DNA repair (Gorbunova et al., 2007; Niedernhofer et al., 2018), and (iv) decreasing performance (aging) at different biological scales (Promislow et al., 2022). Additional empirical support for evolutionary links between SMR and lifespan comes from work showing an inverse cross-species association between SMR and lifespan (Cagan et al., 2022), and from associations between enhanced cancer protection mechanisms and life history traits (MacRae et al., 2015; Tian et al., 2017; Seluanov et al., 2018; Tollis et al., 2020).

The key mediator of model SMR-lifespan coevolution is motivated primarily by cancer for the following reasons: (i) the omnipresence of cancer-associated mutations in certain tissue types, at least for humans (Martincorena, 2019), (ii) the pervasiveness of neoplastic phenomena in multicellular biology (Aktipis et al., 2015; Albuquerque et al., 2018), (iii) evidence that cancers are based on cellular variants, either inherited or acquired (Hanahan, 2022) (but see Frank and Yanai, 2024), (iv) the prevalence of potentially costly mechanisms limiting cancer emergence (Tian et al., 2017; Seluanov et al., 2018; Gorbunova et al., 2014), (v) the temporal separation between mutation events and disease (Gerstung et al., 2020), and (vi) evidence that life history traits can evolve with parasitic diseases (and cancer being analogous in some respects) (Hochberg et al., 1992) and correlate with cancer resistance (Kokko and Hochberg, 2015; Seluanov et al., 2018; Noble et al., 2015).

The impacts of cancers on the evolution of SMR and lifespan are expected to be complex for the following reasons. Similar to the selective pressures regulating cancers occurring before reproductive maturity (Nunney, 2003), post-age-of-reproductive-maturity (but pre-senescent) cancers are expected to exert a negative selective force on both SMR and on lifespan, due to their increased probability of impacting health as a function of age. The balance between evolved defenses that directly target advanced disease versus those focused on preventing neoplastic emergence has received scant attention (Tian et al., 2018). DNA repair and cell variant elimination may expend energy on variants that would not otherwise threaten fitness, whereas mechanisms focused on advanced disease may be energetically costly and less effective once disease is extensive or occurs late in life. Moreover, given the expectation that advanced disease tends to occur at advanced ages (DeGregori et al., 2020; Wright et al., 2003), some of the most health-threatening situations will tend to be partly shielded from natural selection. The late action of disease defenses may be costly or less favored because (i) anti-disease systems age, (ii) disease is increasingly more extensive with age, and (iii) the fitness payoff of defense decreases at sufficiently advanced ages. Depending on the ages at which disease negatively impacts fitness, responses to selection may include some combination of mechanisms influencing SMR (e.g., prevention: DNA mismatch repair, reaction: apoptosis, autophagy) (Kirkwood and Holliday, 1979; Rozhok and DeGregori, 2019), or altered age-specific investment in reproduction (Kirkwood and Shanley, 2005) and homeostasis (Omholt and Kirkwood, 2021) influencing lifespan.

Disease -- and more specifically cancer -- evolution is implicit in the model, insofar as selective disease impacts depend on and modulate both SMR and lifespan. The model assumes continuous evolution (sufficient availability of germline mutant variants), and weak selection on trait means. Because the equations track the evolution of trait means, each right-hand-side term is interpreted as a component of the evolutionary response generated by a selection gradient acting through trait-specific evolvability, in the sense of the canonical equation of adaptive dynamics (Dieckmann and Law, 1996; Geritz et al., 1998). We make this grounding explicit, together with a formal invasion-fitness construction. The model thus follows selective responses only, leaving implicit the following important factors: survival and reproduction (ecology), additive genetic variation and its heritability, distributions in the relative fitnesses of trait variants, epistasis, and trade-offs.

We further assume that SMR and lifespan are each subject to several selective forces that may produce a steady state of trait values, but the mean traits may also be transiently perturbed by environmental vagaries, producing non-equilibrium behaviors. Importantly, the assumption that species traits could equilibrate does not preclude that component selective responses can and will be non-zero at phenotypic trait equilibrium. Finally, as mentioned above, the model focuses on aggregate sources of selection and does not account for details such as the mechanistic roles of specific mutated genes, the precise actions of disease prevention or resistance mechanisms, within-organism mutational or cell dynamics, homeostatic or immune responses, or the age-dependent shape of the force of selection. Accordingly, each parameter should be read as a net, phenomenological selective coefficient rather than as a mechanistically specific quantity.

### Somatic Mutation - Lifespan Coevolution

We present a simple model in which SMR and lifespan coevolve in a single species. These two quantities are encapsulated in a pair of differential equations, 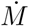 for the evolution of SMR, and 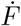 for the evolution of lifespan. The mean population SMR at time *t* is *M*, and the mean lifespan I s *F* .

Table 1 presents definitions of model parameters with *M* and *F* as state variables. We treat lifespan *F*, rather than an instantaneous mortality rate, as the evolving state variable, because *F* is the quantity directly compiled in the comparative data we use (Cagan et al., 2022) and is therefore the life-history state variable used here; it does not encode an age-specific force-of-selection schedule. Mortality is nonetheless represented through the external-hazard parameter *g*. (A fully mortality-based reformulation, in which *F* is replaced by an age-schedule of mortality, is a natural extension; see Discussion). Unless otherwise stated, the constants below are all assumed greater than or equal to zero, before the parameter conventions are applied; after the convention *r* − *l* → *r, r* may be negative.

**Table 1:** Parameters, basic biological interpretation, general entry into model equations, and specific entry at equilibrium, using the parameter conventions *d* − *p* → *d* and *r* − *l* → *r*. See main text for detailed descriptions.

| Parameter | Selection Source | Model Term | Term at Equilibrium |
| --- | --- | --- | --- |
| $Q$ | Body mass | $Qa$ | $Qa$ |
| $a$ | Body-size-scaled selection favoring relaxed somatic maintenance | $Qa$ | $Qa$ |
| $d$ | Disease minus protection/tolerance | $-dMF$ | $-Qa$ |
| $v$ | Body-size-scaled selection favoring somatic function | $Qv$ | $Qv$ |
| $r$ | Net disease/protection effect on lifespan | $-rMF$ | $-rQa/d$ |
| $g$ | External mortality | $-gF$ | $rQa/d - Qv$ |

The model takes the form:

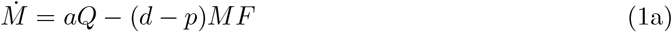

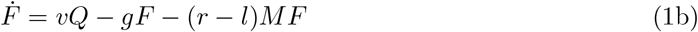

Equations (1a,b) are the response of the resident trait means *z* = (*M, F*) to selection under the canonical equation of adaptive dynamics (Dieckmann and Law, 1996; Geritz et al., 1998),

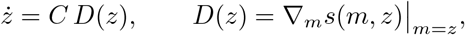

where *s*(*m, z*) is the invasion fitness of a rare mutant with traits (*m, f*) in a resident population (*M, F*), and *D*(*z*) is the corresponding selection gradient. *C* = diag(*c*_*M*_, *c*_*F*_) is a diagonal matrix of positive evolvabilities, i.e., the responsiveness of each trait to selection. We take these as constant, as the basis for the model equations (1a,b).

Each of equations (1a,b) is proportional to the selection gradient on one trait, obtained by asking how the fitness of a rare mutant responds to a small change in that trait. Mutation rate is favored to rise by a body-size-scaled benefit of relaxed somatic maintenance, *aQ*, and is opposed by the fitness cost of disease. That cost grows with the lifetime mutational burden *MF*, because a higher mutation rate produces more lesions per unit time and a longer life allows their consequences to accumulate; its marginal effect on selection against *M* is *dMF* . The net gradient on mutation rate is therefore *aQ* − *dMF*, which is the right-hand side of (1a). By the same reasoning, lifespan is favored by a body-size-scaled benefit *vQ*, reduced by extrinsic mortality *gF*, and modified by the disease-related term *rMF*, giving the gradient *vQ* − *gF* − *rMF*, the right-hand side of (1b).

Because the disease cost acts through the shared product *MF*, the strength of selection on each trait depends on the current value of the other. Selection is thus frequency-dependent and (1a,b) describe the joint coevolution of the two trait means rather than the maximization of a single fixed quantity. This is the expected form for interacting traits under adaptive dynamics (Abrams, 2001).

The model (1a,b) involves two distinct timescales. Physiological (organism) time indexes an individual’s life: the mutation rate *M* is measured in somatic mutations per cell per unit physiological time (e.g., per year), and lifespan *F* is a duration in the same physiological units (e.g., years). Evolutionary (clock) time *t* is the separate, much slower timescale over which the trait means themselves change. The derivatives 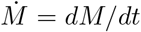 and 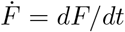 are rates of trait change per unit evolutionary time. The bilinear product *MF* then has the units of a lifetime somatic mutational burden (mutations per cell). The parameters therefore carry mixed units. With body mass *Q* in kg, *a* has units of (mutations per cell per physiological year) per kg per unit evolutionary time; *d*, of inverse (physiological year *×* evolutionary time), so that *d · MF* is a change in *M* per unit evolutionary time; and *g*, of inverse evolutionary time, acting on the physiological duration *F* . We leave these coefficients dimensionally implicit below, noting the two-timescale structure only because it clarifies the phenomenology of the parameters.

*aQ* is a body-size-scaled evolutionary-response term favoring relaxed somatic maintenance (and hence higher *M*), and it is motivated by disposable-soma reasoning (Kirkwood and Rose, 1991; Gomes et al., 2011; Goldsby et al., 2014). Comparative work on mammalian telomeres indicates that larger and/or longer-lived species often evolve altered tumor-suppression architectures, including shorter telomeres (Pepke and Eisenberg, 2022) and, in some clades, altered telomerase activity or proliferative limits as possible protections against cancer (Tian et al., 2018; Risques and Promislow, 2018). We therefore interpret *aQ* not as direct evidence that body size selects for higher somatic mutation rates, but as a phenomenological term capturing body-size-associated selection on the balance between somatic maintenance and other fitness-enhancing investments.

We emphasize that *a* and *v* are not allocation fractions drawn from a shared, fixed budget, and that the model is not age-structured. They are phenomenological, body-size-scaled selection coefficients that enter the invasion fitness (see above) as independent marginal fitness benefits. The disposable-soma framing is therefore used only as a loose motivation for the body-size scaling of these terms, not as an explicit allocation model with a shared budget constraint.

*dMF* is the selective response against SMR due to the downstream fitness costs of disease. Higher mutation rates (*M*) leading to disease can be opposed by enhanced DNA repair (Firsanov et al., 2025), or tolerated since many mutations are benign (Lee-Six et al., 2019; Martincorena et al., 2015). Longer lives (*F*) mean more time for the negative fitness effects of mutation residence and/or mutation accumulation to manifest via disease consequences of, for example, certain somatic mosaicisms or clonal expansions (Cagan et al., 2022). Recent work is consistent with immune-mediated selection and immune evasion during tumor evolution (Lakatos et al., 2025), but this evidence pertains most directly to downstream immune selection or disease-control terms, not to earlier DNA repair unless separate DNA-repair evidence is supplied. For simplicity, we assume that the selective response due to disease is the linear product of SMR (*M*) and lifespan (*F*); that is, marginal increases in lifespan multiply the selective response to lower SMR (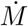) by the current *M* (NB. this is clearly an oversimplification of non-linear growth of cellular expansions; see Discussion). The model is further simplified by assuming the fitness effects of single or small numbers of mutated cells are negligible, and thus a negative selection term proportional to *M* alone can be ignored.

*pMF* is the lowering of selective response against somatic mutation, due to protection against downstream disease. That is, evolved protection acting directly on disease symptoms is assumed to result in the tolerance of mutational effects that would otherwise contribute to disease. Note that protection from downstream disease cannot increase mutation rate beyond the decrease due to disease itself, that is, *p* ≤ *d*.

The evolution of lifespan (eqn. 1b) assumes analogous selective forces to those influencing the evolution of SMR:

*vQ* − *gF* is the net selective response of lifespan independent of somatic mutation. *v* and *g* are constants representing the relative weights of selection based on lifespan increases due to investment in somatic functions (analogous to mechanisms driving *aQ* above) proportional to body mass: *vQ* (Healy et al., 2014), and lifespan decrease due to the cumulative selective effects of external mortality risks, including predators, parasites and abiotic environmental vagaries: *gF* (Kokko and Hochberg, 2015; Hochberg et al., 1992; Johnson et al., 2019; De Vries et al., 2023).

*rMF* is the selective response for shorter lifespan due to downstream disease. Here, rather than reducing SMR through the *dMF* response, another possible response is to shift investments in reproduction to younger ages (e.g., Hochberg et al., 1992), correlatively resulting in earlier ages of senescence and death (Rozhok and DeGregori, 2016). Similar to *dMF*, we assume that the selective response is bilinear; that is, marginal increases in SMR multiply the selective response to lower lifespan (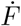) by the current *F* (see Discussion for other functional representations).

Lastly, evolved protection mechanisms that reduce either the incidence or the fitness consequences of downstream somatic pathologies are represented phenomenologically by the term *lMF* (Gorbunova et al., 2014; Tian et al., 2018; Firsanov et al., 2025). Here, *lMF* is intended to capture both direct buffering of pathology at a given burden of somatic lesions, in the sense of tolerance-like protection (Medzhitov et al., 2012; Dillman and Schneider, 2015), and indirect effects whereby protection preserves tissue function and organismal condition. In this latter sense, these indirect effects could alternatively be represented by an increase in effective somatic maintenance, and thus in the positive term *vQ*, consistent with life-history theory in which survival and somatic maintenance are shaped by allocation trade-offs (Kirkwood and Rose, 1991; Maklakov and Chapman, 2019), or by a reduction in the condition-dependent component of realized mortality subsumed in *gF* (De Vries et al., 2023; Chen and Maklakov, 2012). Because *lMF* is defined here to aggregate both direct buffering and these indirect maintenance- or mortality-mediated effects, its net effect need not be bounded by the direct lifespan-shortening effect of pathology, *rMF* ; rather, only the purely direct compensatory component of protection would be expected to oppose *rMF* . We therefore do not impose the constraint *l* ≤ *r* for the aggregate term and instead interpret *lMF* as a first-order approximation to the combined effects of pathology buffering, improved maintenance, and reduced realized mortality risk.

Below, we investigate the two-state equilibrium and its non-equilibrium behavior. Given the same assumed functional forms, in what follows, we use the following parameter conventions for terms in parentheses in eqns. 1a, b: *d*− *p*→ *d* and *r*− *l*→ *r*. The redefined *r* can therefore be negative when protective effects exceed direct disease-mediated lifespan shortening and should be read as a net coefficient rather than as a mortality rate. We refer to *p* and *l* only when needed.

We focus on (i) how parameter values affect equilibrium trait levels and (ii) how a small perturbation to the two-trait equilibrium is met with a non-oscillatory or oscillatory local return to the steady state when the positive interior equilibrium exists. The former (i) considers hypothetical variations in parameter values and resulting relations between equilibrium mutation rate and lifespan. Comparing the shapes of these relations with cross-species data provides only a qualitative consistency check on the underlying model and steady state assumptions. The latter (ii) is intended to represent environmental perturbations that change mean trait levels without changing (evolved) parameter values. There is some empirical evidence for ecological perturbations of genetic variation (trait values) and subsequent recovery (Grant and Grant, 1993; Coulson et al., 2001). Understanding non-equilibrium dynamics and implications for larger-scale phylogeny is important, since it may help explain puzzling empirical observations, for example, evolutionary trajectories that show a time lag or change in direction through time (Avila and Lehmann, 2023; Erten et al., 2025).

### Equilibrium

Equations 1a,b have a single, interior equilibrium point *M* ^∗^, *F* ^∗^ when the positivity conditions below are satisfied and when *a, Q >* 0:

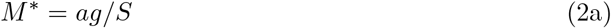

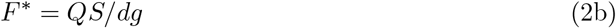

where *S* = *vd* − *ar* represents selective pressure for longer lifespans relative to higher SMR. *S* integrates both intrinsic effects (*a, v*) and interactive effects (*d, r*). At equilibrium, *M* ^∗^ = *ag/S* increases with extrinsic mortality (*g*) and with the net lifespan-shortening coefficient *r*, whereas selection favoring longer lifespans through increased investment in somatic maintenance (*v*) reduces *M* ^∗^, all else being equal. In contrast, equilibrium lifespan *F* ^∗^ = *QS/dg* increases linearly with body mass *Q*, and decreases with extrinsic mortality *g*. Increasing the mutational input parameter *a* reduces *F* ^∗^ when *r >* 0, but increases *F* ^∗^ when *r <* 0. Importantly, although body mass enters the lifespan equilibrium directly, it does not affect equilibrium mutation rate directly when all other parameters are held fixed, a prediction consistent with comparative analyses reporting weak or absent correlations between body mass and somatic mutation rate across species (Cagan et al., 2022). This independence is a direct consequence of the identical linear *Q*-scaling of the two body-size benefit terms, *aQ* and *vQ*, which causes *Q* to cancel from *M* ^∗^. Alternative admissible forms in which the two benefits scale differently with *Q* (e.g., *aQ*^*b*^ for the mutation-rate benefit while the lifespan benefit remains *vQ*, with *b* ≠ 1), or survival-dependent maintenance costs, can induce a body-mass dependence in *M* ^∗^.

As reported in Table 1, the equilibrium selection terms show the pervasiveness of body mass *Q* in arbitrating selective responses in both SMR and lifespan. Most of these terms are dominated by selective responses associated either with costs of genomic maintenance (*a*), or investments in somatic longevity (*v*). Notably, external mortality at equilibrium (− *gF* ^∗^) exactly balances the two other terms in equation (1b), since at equilibrium

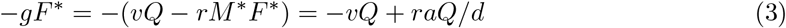

where *M* ^∗^*F* ^∗^ = *aQ/d*. Thus, although *g* enters into both equilibrium levels (2a,b), it does not appear explicitly in the component selective terms listed in Table 1.

The conditions for a positive interior equilibrium point, assuming *a, Q >* 0, are:

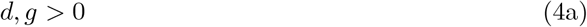

and

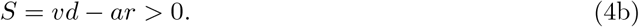

Condition (4a) says that selection against mutation rate due to disease must exceed mutational tolerance resulting from disease protection (in the original parameters, *d > p* ≥ 0 from eq. 1a), and there must be some form of lifespan-limiting mortality (*g >* 0). Condition (4b) says that the product of selection favoring lifespan and selection against mutation, *vd*, must exceed the product of the mutation-promoting term and disease-mediated lifespan shortening, *ar*, when *r >* 0; when *r <* 0, this condition is automatically satisfied for *v, d, a >* 0. When these conditions are violated, *M* or *F* can evolve without bound or to zero (Supplementary Information).

The effects of parameter increases on the interior equilibrium trait levels can be found by taking the partial derivative of *M* ^∗^ = *ag/S* and *F* ^∗^ = *QS/dg*. We have for the effects on equilibrium mutation rate *M* ^∗^:

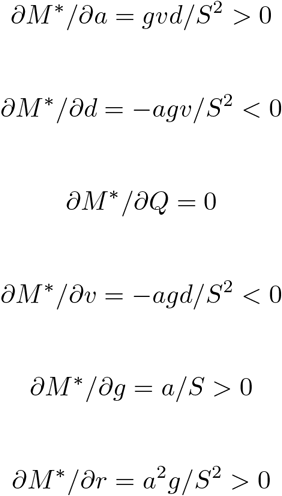

Not surprisingly, selection terms in (1a) translate into parallel effects on equilibrium levels of *M* ^∗^, whereas longer lifespans (high *v*, low *r* in eq. (1b)) decrease equilibrium mutation rate. Note the pervasiveness of parameters *g* and *S*, such that increasing extrinsic mortality *g* increases the magnitude of effects of *a, d, v*, and *r* on *M* ^∗^, whereas -- as is intuitive -- increasing *S* (the net balance defined above) decreases the magnitude of all parameter effects on *M* ^∗^.

Depending on mutational parameters, the impact of mutation may be to increase or decrease equilibrium lifespan *F* ^∗^:

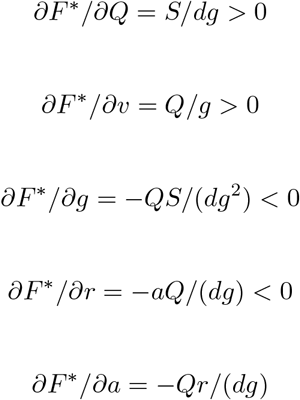

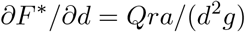

Interestingly, the magnitude of all parameter effects is proportional to body mass *Q* (with the exception of *Q* itself), and inversely proportional to the lifespan-limiting parameter *g*. Interesting too is that the impacts of mutational parameters (*a, d*) on the lifespan equilibrium are contingent on the sign of *r*. Mutation decreases lifespan if *r >* 0, whereas it increases it if *r <* 0. The latter condition corresponds to net protective effects exceeding direct disease-mediated lifespan shortening after the convention *r*− *l*→ *r*. Moreover, if *r <* 0, then a higher mutation-promoting term (high *a*, low *d*) actually increases lifespan.

### Local Stability and Non-Equilibrium Dynamics

We take a first step in understanding the impacts of environmental perturbations on the steady state *M* ^∗^, *F* ^∗^ by considering how small deviations from equilibrium affect the dynamics of *M, F* (see also Supplementary Information). We find that if the interior equilibrium exists, then it is locally stable, but damped oscillatory returns are possible for some *r <* 0 (Figs. 1, 2).

**Figure 1:**
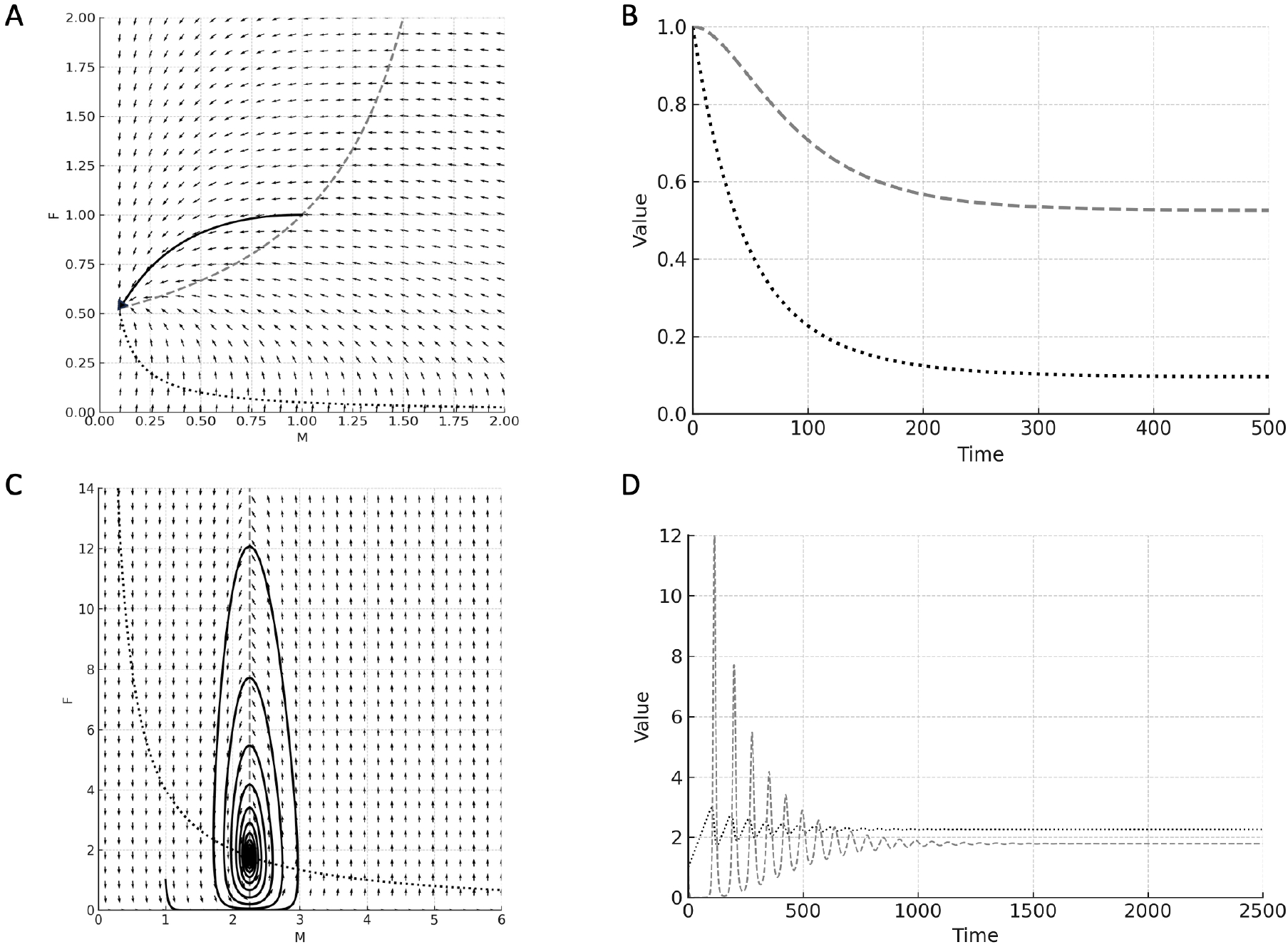
Nullclines and trajectories for the dynamical system. Left panels show phase planes for *M* and *F* ; right panels show time courses of somatic mutation rate *M* and lifespan *F* . In the right panels, *M* is shown as a dotted black curve and *F* as a dashed gray curve. In the left panels, the solid black curve is the phase-plane trajectory starting from initial condition (*M, F*) = (1, 1). Parameter values for A, B are *Q* = 1, *a* = 0.001, *v* = 0.01, *d* = 0.02, *r* = − 0.01, and *g* = 0.02, giving a stable-node equilibrium at approximately (*M* ^∗^, *F* ^∗^) = (0.095, 0.525). Parameter values for C, D are *Q* = 1.0, *a* = 0.02, *v* = 0.0001, *d* = 0.005, *r* = − 0.4, and *g* = 0.9, giving a stable-focus equilibrium at approximately (*M* ^∗^, *F* ^∗^) = (2.250, 1.778).

**Figure 2:**
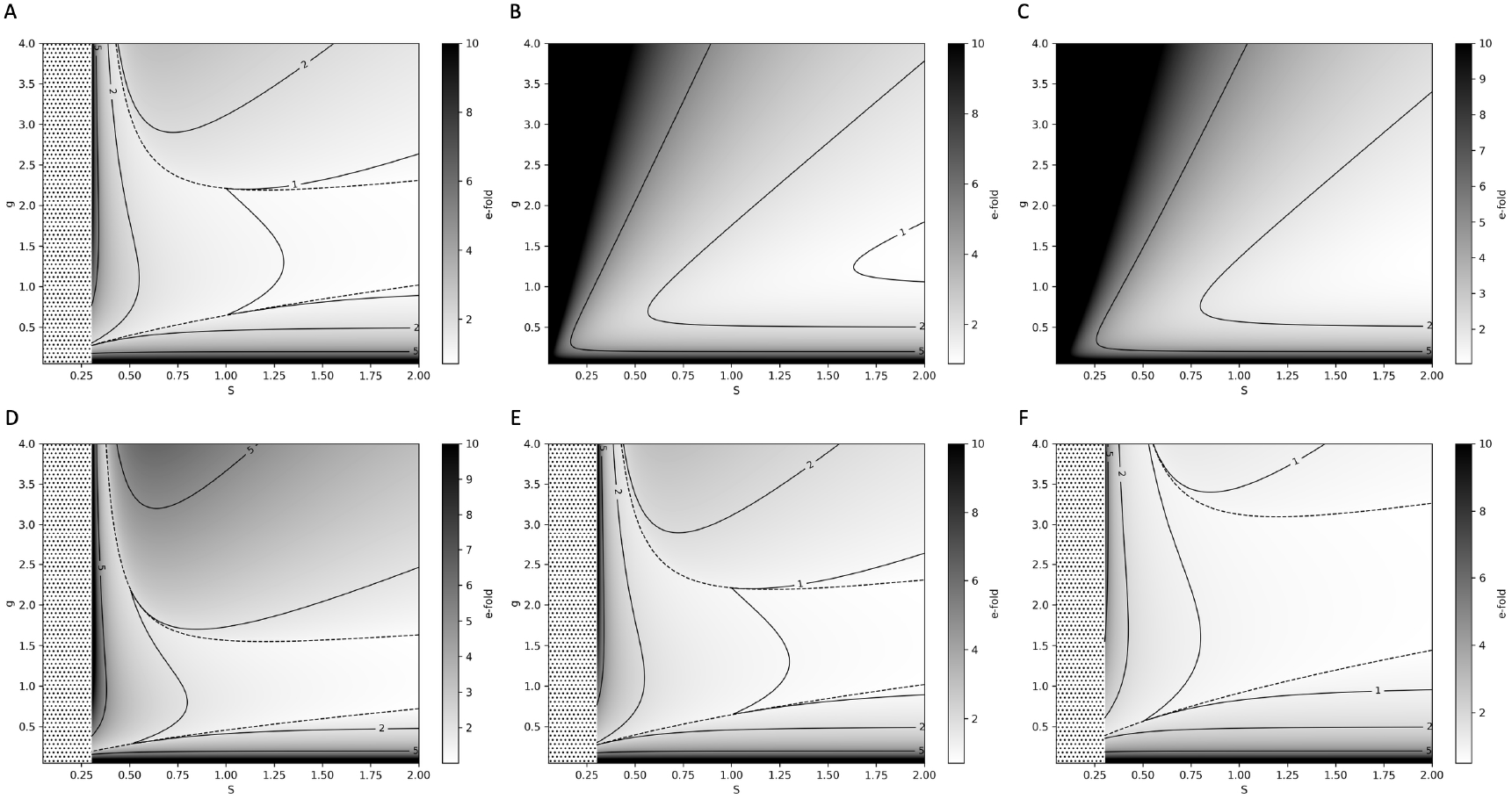
E-fold return time to the interior equilibrium *M* ^∗^, *F* ^∗^ following a small perturbation. E-fold values are truncated at 10. Darker shading corresponds to slower returns. Solid black curves are contours of e-fold return time. Dashed curves in A, D, E, and F delimit the stable-focus region, corresponding to damped oscillatory return, from the stable-node region, corresponding to non-oscillatory return. Stippled areas in A and D–F indicate parameter values inadmissible under the assumption *v >* 0: with *a* = *d* = 1 and *r* = − 0.3, these values would require *v* = *S* − 0.3 ≤ 0. They should not be interpreted as regions with no interior equilibrium. The plotted axes are *S* and *g*. Fixed parameters not otherwise specified are set to 1.0, with *v* varied according to *v* = *S* + *r* to obtain the plotted values of *S*. A: *r* = −0.3, *Q* = 1.0, *v* ∈ (0, 1.7]; B: *r* = 0.1, *Q* = 1.0, *v* ∈ [0.15, 2.1]; C: *r* = 0.3, *Q* = 1.0, *v* ∈ [0.35, 2.3]; D: *r* = −0.3, *Q* = 0.5, *v* ∈ (0, 1.7]; E: *r* = −0.3, *Q* = 1.0, *v* ∈ (0, 1.7]; F: *r* = −0.3, *Q* = 2.0, *v* ∈ (0, 1.7]. Panels A and E are identical and are repeated for comparative purposes.

Local stability of the interior equilibrium requires

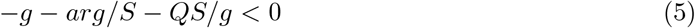

equivalently, using *S* + *ar* = *vd*,

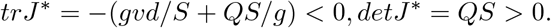

These conditions are always satisfied whenever the interior equilibrium exists. Furthermore, the equilibrium is a locally stable focus (damped oscillatory return) if and only if

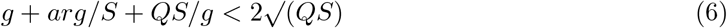

a condition that can only be met when *S >* 0 and for some *r <* 0; otherwise, the equilibrium is a locally stable node (non-oscillatory local returns to equilibrium). Visual inspection of Fig. 2 indicates that the condition for damped oscillations holds for intermediate parameter values (Supplementary Information).

Linearized return times to the interior equilibrium, quantified by an e-fold index (Strogatz, 2018), provide a standard local measure of non-equilibrium behavior near a steady state. The parameter effects are complex (Fig. 2; Supplementary Information). First, in the stable-focus regime (damped oscillations) of the *M* ^∗^, *F* ^∗^ equilibrium, increasing body mass *Q* decreases return times, thereby accelerating return. Moreover, within the focus regime, increasing either *v* or *d* also decreases return times, because both parameters strengthen the stabilizing feedbacks that govern decay of small perturbations. In contrast, the effects of *a, r*, and *g* in the focus regime are contingent on the sign of the expression *QS*^2^ − *dg*^2^*v*, reflecting the relative levels of coupling strength (bilinear *MF* terms) and dissipation (*g*). Second, in the stable-node regime, e-fold return times depend jointly on how parameters shift stabilizing versus destabilizing effects. Changes in *a, v*, and *d* can simultaneously strengthen some stabilizing contributions while weakening others, reducing the scope for general monotonic predictions of how e-fold times vary with parameters.

### Analysis of Cross-Species Patterns

Evaluating the theory developed here will require data from either (i) published accounts or controlled experiments tracking the evolution of single-species state variables under different conditions, or (ii) cross-species comparisons of variables corresponding to parameters and state variables in equations 1a,b. We are unaware of data for animals to accomplish (i). In contrast, data from a recent study (Cagan et al., 2022) can be employed to evaluate expected relations, based on equilibrium assumptions.

Cagan and colleagues generated somatic mutation-rate estimates from intestinal crypt sequencing across 16 mammalian species; age-informed analyses were possible for 15 species, and the study also compiled lifespan and body-mass estimates. The data are limited in numerous ways, most importantly in being phylogenetically restricted, based on small sample sizes, and reporting measures at the species level (e.g., not considering different breeds of dogs). As a first step, we employ their mean estimates, equating their somatic mutation rate with our equilibrium SMR, *M* ^∗^, their lifespan with our equilibrium lifespan, *F* ^∗^, and their body mass with the appearance of *Q* in the equilibrium of lifespan. These empirical lifespans are proxies for *F* ^∗^; they are not direct estimates of the age at which the demographic force of selection becomes negligible. Because the species sample is small and phylogenetically restricted, we do not treat their *p*-values as confirmatory tests of the model; a broader phylogenetically corrected analysis would be necessary to evaluate our findings.

Cagan and colleagues found a strong inverse relationship between lifespan and mutation rate, which appears convex on untransformed axes. Our equilibrium relations (equations 2a,b) can approximate this relationship parsimoniously if *S*→ 0^+^, yielding high mutation rate and short lifespan, and *g*→ 0^+^, yielding low mutation rate and long lifespan. The former condition corresponds to the limit of equilibrium validity, whereas the latter says that low selection from external mortality alone can drive low mutation rate and high lifespan. This is because of the action of *g* in mediating coevolution: low *g* results in longer lifespans, and the bilinear disease term then requires lower SMR at equilibrium so that *M* ^∗^*F* ^∗^ = *aQ/d*. Also possible is that the empirical shape of the lifespan-mutation rate curve could be explained by co-variation in model parameters (e.g., high *Q*, high *S*, giving low SMR and long lifespan; low *Q*, low *S*, giving high SMR and short lifespan).

We then examined two derived measures: expected mutations accumulated over a lifespan, *M* ^∗^*F* ^∗^ (Fig. 3A), and SMR normalized by lifespan, *M* ^∗^*/F* ^∗^ (Fig. 3B). In both assessments we assume that lifespan in our model is approximated by the species-level lifespan estimates reported by (Cagan et al., 2022). Note that *M* ^∗^*/F* ^∗^ is not a lifetime burden: if *M* ^∗^ is a per-year mutation rate, *M* ^∗^*/F* ^∗^ has units of mutation rate divided by time (e.g., mutations per cell per year squared) and is best treated as an exploratory contrast derived from the model, not as a directly observed biological quantity.

**Figure 3:**
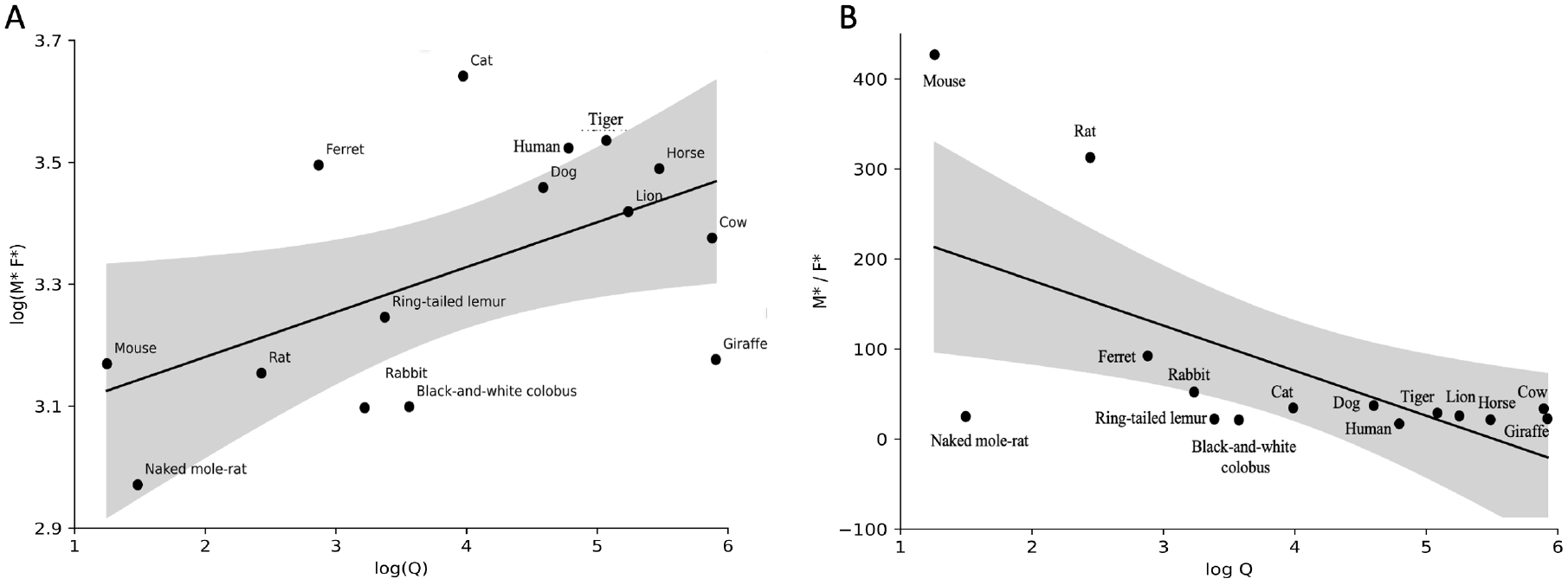
A. Regression of log *Q* vs log *M* ^∗^*F* ^∗^ for data from 15 species given in Cagan et al. 2022. *R*^2^ = 0.298, *p* = 0.035. B. Regression of log *Q* vs *M* ^∗^*/F* ^∗^ for data from 15 species given in Cagan et al. 2022. *R*^2^ = 0.387, *p* = 0.0175. Shaded area is 95% confidence limits based on Student’s *t*, df = 13. Panel B should be interpreted as an exploratory regression of a derived index rather than as a direct mutation-burden measure.

Figure 3A plots how log *Q* associates with log *M* ^∗^*F* ^∗^, the latter being a measure of mutation accumulation over a lifespan, and predicted by our model, at equilibrium, to be *aQ/d*, a quantity with units of lifetime mutational burden under linear accumulation independent of the lifespan-specific parameters *g* and *S*. A proportional relationship between *M* ^∗^*F* ^∗^ and *Q* across species therefore requires *a/d* to be constant; because Cagan et al. reported approximate conservation of end-of-lifespan burden despite orders-of-magnitude variation in body mass, the model would require *a/d* to decrease approximately as 1*/Q*, or the body-mass term *Q* to be interpreted more narrowly than literal adult mass. The *MF* term is phenomenological and should not be identified with a mechanistically defined latent burden. The empirical product used in Fig. 3A is computed from the observed mutation-rate and lifespan estimates; any washing-out effect therefore remains a separate hypothesis (Fig. 4B), not an explanation supplied by the equilibrium model. Based on previous analyses, we expect little or no relationship between body mass *Q* and SMR (Cagan et al., 2022; Vincze et al., 2022). However, when considering mutations over a lifespan (log *M* ^∗^*F* ^∗^) the non-phylogenetic regression is nominally significant with a positive slope, suggesting that, in this dataset, larger-bodied species may have higher estimated lifetime mutational burden than smaller-bodied species. This pattern is consistent with, but does not by itself demonstrate, increased protection or tolerance of somatic mutation in high-body-mass species.

**Figure 4:**
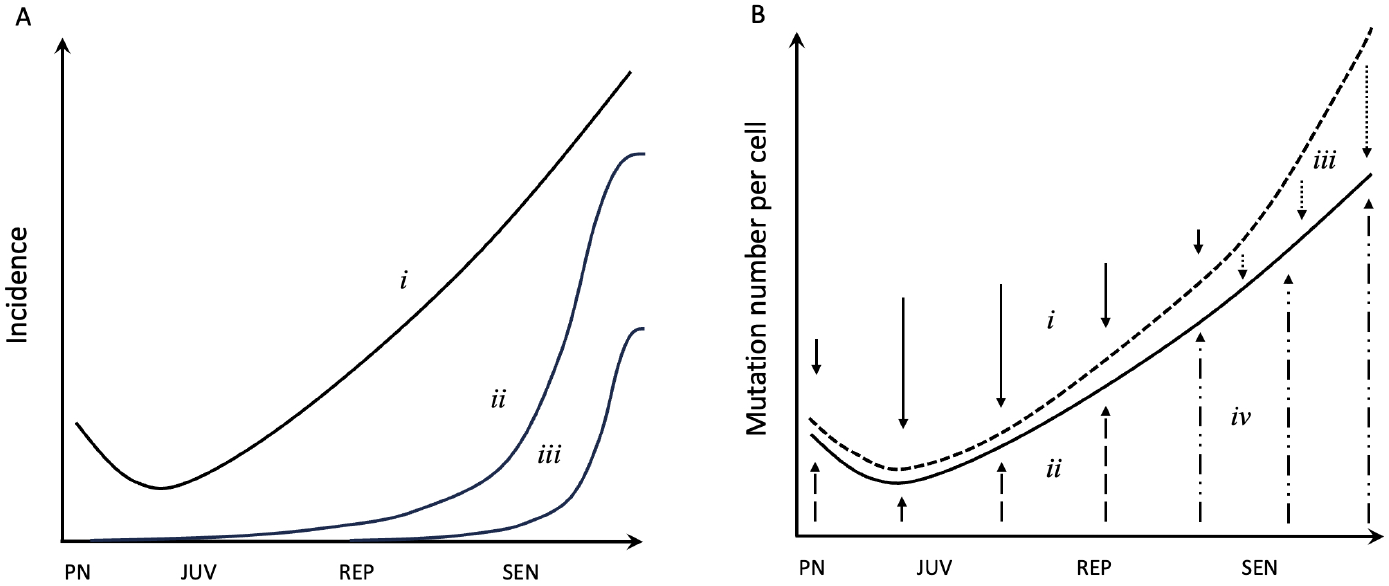
Hypothetical trends in key measures and events. A. i: schematic shape of observed mutation accumulation through the lifespan; ii: hypothetical wall of disease; iii: wall of death. B. Hypothesis for the possible non-detection of exponential increases in mutations with age. Lower curve: observed; upper curve: actual. i: selection for protection against disease; ii: tolerance of early mutations; iii: mutation loads never observed because the most highly mutated cells are lost earlier; iv: tolerance of late mutations. See main text for details. Abbreviations: PN = prenatal; JUV = juvenile; REP = reproductive; SEN = senescence.

In contrast, when plotting the data as log *Q* vs. SMR normalized by lifespan, *M* ^∗^*/F* ^∗^, we see that the regression slope is negative, such that larger-bodied organisms have lower lifespan-normalized SMR (Fig. 3B). Cagan and coworkers found that mutation rate was strongly inversely associated with lifespan and only weakly associated with body mass before, and not after, accounting for lifespan; they also reported approximate conservation of end-of-lifespan burden across species (Cagan et al., 2022). Note that the negative slope in our regression is disproportionately due to short-lived, small body-sized mice and rats, whose removal results in an approximately flat relationship (see also Zhang et al., 2021; Cagan et al., 2022). More broadly, because these are cross-species correlations, they cannot by themselves establish the direction of causation: the inverse somatic mutation rate–lifespan relationship is equally compatible with a lower somatic mutation rate promoting longer life, with longer life selecting for stronger genome maintenance, or with both traits being shaped by shared life-history or mechanistic determinants (and cancer risk being a cause, a consequence, or a correlate of these). We therefore treat the disease-mediated coevolution modeled here as one hypothesis consistent with these data, not as a causal inference drawn from the correlations.

## Discussion

Understanding mutation in both the germline and in the soma is a central challenge in disease biology and more generally in evolutionary biology. Somatic genomes and cellular organization are continually challenged by damage and error, and maintaining function requires protective and repair processes (Vijg, 2021; Schumacher et al., 2021; Eigen, 1971; Szathmáry, 2015). In this sense, SMR may represent a potentially fundamental constraint on the evolution of lifespan, because longer lives increase the time over which somatic informational loss can accumulate and manifest as disease, unless offset by genome maintenance, tissue-level control, or tolerance (Cagan et al., 2022; Gorbunova et al., 2014).

Empirical study reveals the daunting complexity of mutational processes and the factors responsible for variations within individuals, between individuals in a population, and between species. Beyond the necessity that comparative and experimental approaches seek to uncover the inner workings of this complexity, mathematical theory has an important role to play in exploring possible interrelations and predicted patterns, which can, in turn, form the basis for empirical tests. Although we view cancer as one candidate driver of the patterns elucidated here, the model concepts are general to any condition or disease resulting from somatic mutation or other breakdowns in homeostasis, threatening to grow in severity with age, impacting individual fitness, and susceptible to the evolution of prevention or remediation.

## Main results

The model yields four main results. First, we found that approaching the limit conditions for a positive equilibrium is associated either with a linear decrease in lifespan and a nonlinear increase in somatic mutation rate (SMR), or the reverse. This is qualitatively consistent with comparative findings (Cagan et al., 2022). Our model suggests that the overall shape of the relationship can be parsimoniously explained by the selective pressure for longer lifespans *S* and the direct impact of external mortality (*g*) on lifespan. Specifically, the non-linear effect is driven by low *g* in low-SMR, long-lifespan species and by *S*→ 0 in high-SMR, short-lifespan species. The same pattern can also emerge if body mass (*Q*) co-varies across species with *S*, the net selection favoring longer lifespan over higher mutation rate: large *Q* combined with high *S* yields low SMR and long lifespan, whereas small *Q* with low *S* yields the reverse. Second, under the specified identical linear body-size scaling, we found that body size (*Q*) associates with equilibrium lifespan, but not SMR, when other model coefficients are held fixed, again, consistent with empirical findings (Cagan et al., 2022). Although not considered in our model, over sufficiently long timescales we would expect body size (*Q*) to evolve (Kokko and Hochberg, 2015). Should selection over long timescales reduce body size, then our model would predict no change in SMR at equilibrium only if the remaining parameters do not covary with body size, and given the observed positive association between SMR and neoplasms in a limited comparative sample (Compton et al., 2025), this would be consistent with recent theoretical findings suggesting that evolutionary decreases in body size do not necessarily result in increases in cancer prevalence (Erten et al., 2020, 2025). Third, we showed that a positive trait equilibrium can only be achieved if *S* = *vd* − *ar >* 0.

Finally, damped oscillatory returns are possible only when the net lifespan coefficient is negative (*r <* 0 after the parameter convention), and then only when condition (6) is satisfied. This effect could arise, for example, if reduced disease symptoms result in reduced predation or parasitism (Hochberg and Noble, 2017). Some long-lived species show evidence for defense mechanisms against mutation accumulation and cancer (Zhang et al., 2021; Tian et al., 2017; Seluanov et al., 2018; Hochberg et al., 2016), but it is not known to what extent reduced external mortality, as opposed to reduced mortality due to cancer itself, contributes to selection for longer lifespans.

Comparative studies of somatic mutation rate, lifespan, body size, and cancer risk do not by themselves establish whether species are at trait equilibrium (Cagan et al., 2022; Vincze et al., 2022). The present study considers the more general situation in which traits may or may not be at equilibrium, and provides predictions both for trait levels and for their local evolutionary trajectories. Natural populations undergoing recent environmental change, and confined populations experiencing altered mortality, reproduction, veterinary care, or disease exposure, therefore provide useful testing grounds for the theory presented here.

### A hypothesis concerning age-related mutation accumulation

Should pathologies be prevented, attenuated, or delayed by selection (Kokko and Hochberg, 2015; Nunney, 2003; Frank, 2007), then theory would suggest that many mutation-linked or age-related maladies should increasingly parallel the inverse of the force of selection, that is, tend to accelerate as selection wanes. If this is the case, then the selection logic motivating our model suggests a hypothesis in which tolerated mutations and, more generally, somatic anomalies, accumulate and contribute to aging phenotypes (Chatsirisupachai and De Magalhães, 2024), while shifting part of disease incidence toward ages at which natural selection is weak. Available data are consistent with late-life nonlinear increases in age-related disease burden (Marongiu and DeGregori, 2022; Shen et al., 2024), potentially resembling a hypothetical “wall of disease” at ages where selection is greatly reduced, but not zero (for cancer see Rozhok and DeGregori, 2020, 2016; Nunney, 2003) (Fig. 4A). Contrast this with the “wall of death”, which can arise in some mutation-accumulation formulations when survivorship collapses as the force of selection approaches zero, but is not a universal prediction of evolutionary aging theory (Charlesworth, 2001). Overlapping disease incidences could be a consequence of selection as posited here, but might also result from aging-associated tissue change and clonal selection (Marongiu and DeGregori, 2022), or, more simply, from similar rates of disease development independent of aging. To the extent that certain diseases and conditions are a product of single mutations (somatic mosaicism), breakdown in homeostasis (aging), or clonally expanding cells (cancer), specific predictions of appropriately modified models should be possible concerning age-incidence profiles.

Consistent with theory on the force of selection, DNA repair can decline with age in mammals (Gorbunova et al., 2007), suggesting that somatic aberrations should accumulate with age. Considerable data are consistent with an approximately linear increase in mutation load (e.g., Cagan et al., 2022; Alexandrov et al., 2015; Blokzijl et al., 2016; Moore et al., 2021; Osorio et al., 2018; Abascal et al., 2021), although some work has proposed nonlinear or even exponential increases (Milholland et al., 2017). This discrepancy may be real, or it may reflect some combination of technical limitations and unaccounted-for biological processes. In particular, somatic mutational burden in normal tissues is not easily quantified and may therefore be underestimated (Chatsirisupachai and De Magalhães, 2024). However, it is not straightforward to see why such effects would disproportionately manifest at older ages. By contrast, evolved mutational tolerance, together with the subsequent “washing out” of highly mutated cells through differentiation, senescence, or cell death (Nowak et al., 2003; Derényi and Szöllősi, 2017; Demeter et al., 2022), speculatively offers a possible mechanistic explanation for why exponential increases may not be observed (Fig. 4B). Although our model makes no specific predictions regarding age profiles of mutation accumulation, it does provide the parameter relations that modulate SMR normalized by lifespan (as a derived rate-lifespan contrast, not a direct mutation burden): *adg*^2^*/*(*QS*^2^). Higher lifespan-normalized SMR should occur with some combination of low body mass (*Q*), high external mortality (*g*), and a close balance in selection pressure between the competing demands of somatic protection and somatic growth (low *S*). This broader expectation that short-lived species should experience higher per-time somatic mutation rates is consistent with comparative findings that somatic mutation rates scale inversely with lifespan, and that some long-lived species show evidence of enhanced protective mechanisms (Cagan et al., 2022; Gorbunova et al., 2014).

## Limitations

All else being equal, somatic mutations arising early in development can be propagated to a larger fraction of the body and therefore carry greater pathogenic potential than mutations arising later in life (Manders et al., 2021; Kuijk et al., 2019; Youssoufian and Pyeritz, 2002). Early driver mutations may thus increase the number of cells susceptible to subsequent malignant progression, and some cancer-driving mutations appear to originate years to decades before diagnosis (Gerstung et al., 2020; Beerenwinkel et al., 2007). The models presented here do not explicitly incorporate age-specific processes and therefore are not intended to generate detailed ontogenetic predictions. Nevertheless, available human data are consistent with higher prenatal mutation rates in some tissues and developmental contexts; claims about relaxed genome surveillance, cell-cycle checkpoints, or apoptosis require mechanism-specific evidence beyond elevated mutation rates alone, although these findings do not by themselves demonstrate relaxed evolutionary selection, or mutational tolerance in the sense used here (Manders et al., 2021; Kuijk et al., 2019). This timing matters because progression from an initiating mutation to life-threatening cancer often requires prolonged clonal expansion and additional hits, whereas the force of natural selection is generally strongest earlier in life and declines with age (Gerstung et al., 2020; Johnson et al., 2019; Beerenwinkel et al., 2007). Somatic changes that arise early enough to influence survival or reproduction across a substantial portion of the reproductive lifespan should therefore be more strongly exposed to negative selection than equivalent changes whose effects are confined largely to late life. Overall, these considerations support the view that both the costs of maintaining cellular integrity and the downstream fitness consequences of disease can shape life-history evolution, including lifespan.

A further limitation concerns our focus on the evolution of somatic mutation rate only. Germline and somatic mutation rates are correlated across species and are expected to share mechanistic determinants, including DNA-repair and replication fidelity, metabolic rate, generation time, and overall life-history pace (Moore et al., 2021; Bergeron et al., 2023). Selection on germline mutation rate can act through mutational effects transmitted to descendants, whereas selection on somatic mutation rate acts through effects on the carrier’s survival and reproduction; their relative strengths are context dependent (Lynch, 2010a; Lynch et al., 2016). Consequently, part of the cross-species association between somatic mutation rate and lifespan analyzed here could be a correlated by-product of selection on germline mutation rate, or of shared determinants of both rates, rather than specifically requiring disease-mediated coevolution of somatic mutation rate only.

### Extensions and future research

Although the theory developed here focuses on cancer, we suggest it applies more generally to age-dependent diseases and conditions. This includes clonal expansions other than cancer (Hall et al., 2019; Jaiswal and Ebert, 2019) and somatic mosaicism (De, 2011), but possibly also diseases associated with aging (Guo et al., 2022) and perhaps aging itself (Promislow et al., 2022; Li et al., 2015; Pomatto and Davies, 2017). For our model to apply to aging, associated fitness deficits must be subject to a selective response (i.e., heritable genetic variation for prevention or remediation). More generally, specific diseases or conditions may have characteristic age-dependent dynamics, meaning that selective responses can target early intracellular disruption (e.g., DNA repair, proteostasis), subsequent cellular or tissue functioning defects (e.g., apoptosis, autophagy, immune responses, senescence), or larger-scale life history (e.g., development, reproductive effort, body size). Untangling the roles of somatic mutation and costs associated with evolved protection in the aging process will be highly challenging (Vijg, 2021; Schumacher et al., 2021; Vijg and Dong, 2020; Kennedy et al., 2012; De Magalhães, 2025).

Future research should consider: (1) Greater theoretical integration of ecological and evolutionary dynamics, particularly to understand how short-term demographic processes interact with long-term selection on somatic maintenance and life-history traits. (2) More explicit quantitative-genetic or population-genetic models with standing variation, covariance, and genetic architecture, building on the selection-gradient formulation introduced here toward fully dynamical quantitative-genetic or population-genetic treatments. (3) Age-specific schedules of both external mortality and trait expression, as selective pressures and phenotypic effects are unlikely to be constant across the lifespan. Relatedly, recasting the lifespan state variable in terms of an explicit age-schedule of mortality would provide more explicit demographic building blocks than simple lifespan. (4) Tissue-specific differences in the coevolution of somatic mutation rates and life-history traits. (5) Mutation rate-lifespan coevolution within a broader multivariate framework that includes other key traits, such as body size and reproductive effort. (6) Models tailored to specific age-dependent diseases and pathological conditions, enabling closer connections between evolutionary predictions and biomedical data. (7) Coevolutionary models that include germline mutation processes, explicitly allowing somatic mutation rate to evolve as a correlated response to selection on germline mutation rate, and to shared mechanistic determinants of both rates. (8) Comparative phylogenetic analyses focusing on variation in evolutionary rates, with particular emphasis on identifying and explaining lineages that deviate from general scaling relationships. (9) How interspecific variation in regressions linking cancer incidence and life-history parameters may reflect the evolution of enhanced cancer protection mechanisms, increased tolerance of somatic mutation, or both.

## Acknowledgments

I thank numerous colleagues for discussions, and acknowledge generous core support from my host institutions, and funding from the Fondation ARC pour la recherche sur le cancer: Programmes labellisés 2021, Project ARCPGA12021010002850 3574.

## References

Abascal, F., Harvey, L. M. R., Mitchell, E., Lawson, A. R. J., Lensing, S. V., Ellis, P., Russell, A. J. C., Alcantara, R. E., Baez-Ortega, A., Wang, Y., Kwa, E. J., Lee-Six, H., Cagan, A., Coorens, T. H. H., Chapman, M. S., Olafsson, S., Leonard, S., Jones, D., Machado, H. E., Davies, M., Øbro, N. F., Mahubani, K. T., Allinson, K., Gerstung, M., Saeb-Parsy, K., Kent, D. G., Laurenti, E., Stratton, M. R., Rahbari, R., Campbell, P. J., Osborne, R. J., and Martincorena, I. (2021). Somatic mutation landscapes at single-molecule resolution. Nature, 593(7859):405–410.

Abrams, P. A. (2001). Modelling the adaptive dynamics of traits involved in inter- and intraspecific interactions: An assessment of three methods. Ecology Letters, 4(2):166–175.

Aktipis, C. A., Boddy, A. M., Jansen, G., Hibner, U., Hochberg, M. E., Maley, C. C., and Wilkinson,G. S. (2015). Cancer across the tree of life: Cooperation and cheating in multicellularity. Philosophical Transactions of the Royal Society B: Biological Sciences, 370(1673):20140219.

Albuquerque, T. A. F., Drummond Do Val, L., Doherty, A., and De Magalhães, J. P. (2018). From humans to hydra: Patterns of cancer across the tree of life. Biological Reviews, 93(3):1715–1734.

Alexandrov, L. B., Jones, P. H., Wedge, D. C., Sale, J. E., Campbell, P. J., Nik-Zainal, S., and Stratton, M. R. (2015). Clock-like mutational processes in human somatic cells. Nature Genetics, 47(12):1402–1407.

Avila, P. and Lehmann, L. (2023). Life history and deleterious mutation rate coevolution. Journal of Theoretical Biology, 573:111598.

Baudisch, A. (2005). Hamilton’s indicators of the force of selection. Proceedings of the National Academy of Sciences, 102(23):8263–8268.

Beerenwinkel, N., Antal, T., Dingli, D., Traulsen, A., Kinzler, K. W., Velculescu, V. E., Vogelstein, B., and Nowak, M. A. (2007). Genetic Progression and the Waiting Time to Cancer. PLoS Computational Biology, 3(11):e225.

Bergeron, L. A., Besenbacher, S., Zhang, J., et al. (2023). Evolution of the germline mutation rate across vertebrates. Nature, 615(7951):285–291.

Blokzijl, F., De Ligt, J., Jager, M., Sasselli, V., Roerink, S., Sasaki, N., Huch, M., Boymans, S., Kuijk, E., Prins, P., Nijman, I. J., Martincorena, I., Mokry, M., Wiegerinck, C. L., Middendorp, S., Sato, T., Schwank, G., Nieuwenhuis, E. E. S., Verstegen, M. M. A., Van Der Laan, L. J. W., De Jonge, J., IJzermans, J. N. M., Vries, R. G., Van De Wetering, M., Stratton, M. R., Clevers, H., Cuppen, E., and Van Boxtel, R. (2016). Tissue-specific mutation accumulation in human adult stem cells during life. Nature, 538(7624):260–264.

Bowling, S., Lawlor, K., and Rodríguez, T. A. (2019). Cell competition: The winners and losers of fitness selection. Development, 146(13):dev167486.

Buckingham, L. J. and Ashby, B. (2024). Coevolution of age-structured tolerance and virulence. Bulletin of Mathematical Biology, 86(6):62.

Cagan, A., Baez-Ortega, A., Brzozowska, N., Abascal, F., Coorens, T. H. H., Sanders, M. A., Lawson, A.R. J., Harvey, L. M. R., Bhosle, S., Jones, D., Alcantara, R. E., Butler, T. M., Hooks, Y., Roberts, K., Anderson, E., Lunn, S., Flach, E., Spiro, S., Januszczak, I., Wrigglesworth, E., Jenkins, H., Dallas, T., Masters, N., Perkins, M. W., Deaville, R., Druce, M., Bogeska, R., Milsom, M. D., Neumann, B., Gorman, F., Constantino-Casas, F., Peachey, L., Bochynska, D., Smith, E. S. J., Gerstung, M.,Campbell, P. J., Murchison, E. P., Stratton, M. R., and Martincorena, I. (2022). Somatic mutation rates scale with lifespan across mammals. Nature, 604(7906):517–524.

Cairns, J. (1975). Mutation selection and the natural history of cancer. Nature, 255(5505):197–200.

Campisi, J. (2005). Aging, tumor suppression and cancer: High wire-act! Mechanisms of Ageing and Development, 126(1):51–58.

Charlesworth, B. (2001). Patterns of Age-specific Means and Genetic Variances of Mortality Rates Predicted by the Mutation-Accumulation Theory of Ageing. Journal of Theoretical Biology, 210(1):47–65.

Chatsirisupachai, K. and De Magalhães, J. P. (2024). Somatic mutations in human ageing: New insights from DNA sequencing and inherited mutations. Ageing Research Reviews, 96:102268.

Chen, H.-y. and Maklakov, A. A. (2012). Longer life span evolves under high rates of condition-dependent mortality. Current Biology, 22(22):2140–2143.

Cheng, C. and Kirkpatrick, M. (2021). Molecular evolution and the decline of purifying selection with age. Nature Communications, 12(1):2657.

Colom, B., Herms, A., Hall, M. W. J., Dentro, S. C., King, C., Sood, R. K., Alcolea, M. P., Piedrafita, G., Fernandez-Antoran, D., Ong, S. H., Fowler, J. C., Mahbubani, K. T., Saeb-Parsy, K., Gerstung, M., Hall, B. A., and Jones, P. H. (2021). Mutant clones in normal epithelium outcompete and eliminate emerging tumours. Nature, 598(7881):510–514.

Compton, Z. T., Mellon, W., Harris, V. K., Rupp, S., Mallo, D., Kapsetaki, S. E., Wilmot, M., Kennington, R., Noble, K., Baciu, C., Ramirez, L. N., Peraza, A., Martins, B., Sudhakar, S., Aksoy, S., Furukawa, G., Vincze, O., Giraudeau, M., Duke, E. G., Spiro, S., Flach, E., Davidson, H., Li, C. I., Zehnder, A., Graham, T. A., Troan, B. V., Harrison, T. M., Tollis, M., Schiffman, J. D., Aktipis, C. A., Abegglen, L. M., Maley, C. C., and Boddy, A. M. (2025). Cancer Prevalence across Vertebrates. Cancer Discovery, 15(1):227–244.

Coulson, T., Catchpole, E. A., Albon, S. D., Morgan, B. J. T., Pemberton, J. M., Clutton-Brock, T. H., Crawley, M. J., and Grenfell, B. T. (2001). Age, sex, density, winter weather, and population crashes in soay sheep. Science, 292(5521):1528–1531.

Dang, C. V. (2015). A metabolic perspective of Peto’s paradox and cancer. Philosophical Transactions of the Royal Society B: Biological Sciences, 370(1673):20140223.

De, S. (2011). Somatic mosaicism in healthy human tissues. Trends in Genetics, 27(6):217–223.

De Magalhães, J. P. (2025). The evolution of cancer and ageing: A history of constraint. Nature Reviews Cancer, 25(11):873–880.

De Vries, C., Galipaud, M., and Kokko, H. (2023). Extrinsic mortality and senescence: A guide for the perplexed. Peer Community Journal, 3:e29.

DeGregori, J., Pharoah, P., Sasieni, P., and Swanton, C. (2020). Cancer Screening, Surrogates of Survival, and the Soma. Cancer Cell, 38(4):433–437.

Demeter, M., Derényi, I., and Szöllősi, G. J. (2022). Trade-off between reducing mutational accumulation and increasing commitment to differentiation determines tissue organization. Nature Communications, 13(1):1666.

Derényi, I. and Szöllősi, G. J. (2017). Hierarchical tissue organization as a general mechanism to limit the accumulation of somatic mutations. Nature Communications, 8:14545.

D’Gama, A. M. and Walsh, C. A. (2018). Somatic mosaicism and neurodevelopmental disease. Nature Neuroscience, 21(11):1504–1514.

Dieckmann, U. and Law, R. (1996). The dynamical theory of coevolution: A derivation from stochastic ecological processes. Journal of Mathematical Biology, 34(5-6):579–612.

Dillman, A. R. and Schneider, D. S. (2015). Defining resistance and tolerance to cancer. Cell Reports, 13(5):884–887.

Eigen, M. (1971). Selforganization of matter and the evolution of biological macromolecules. Die Naturwissenschaften, 58(10):465–523.

Erten, E. Y., Tollis, M., and Kokko, H. (2020). Bird size with dinosaur-level cancer defences: Can evolutionary lags during miniaturisation explain cancer robustness in birds? BioRxiv [Preprint].

Erten, E. Y., Tollis, M., and Kokko, H. (2025). Shrinking to bird size with dinosaur-level cancer defences: Evolution of cancer suppression over macroevolutionary time. PLOS Computational Biology, 21(9):e1013432.

Evans, E. J. and DeGregori, J. (2021). Cells with cancer-associated mutations overtake our tissues as we age. Aging and Cancer, 2(3):82–97.

Evans, M. A. and Walsh, K. (2023). Clonal hematopoiesis, somatic mosaicism, and age-associated disease. Physiological Reviews, 103(1):649–716.

Failla, G. (1958). The Aging Process and Cancerogenesis. Annals of the New York Academy of Sciences, 71(6 Genetic Conce):1124–1140.

Fane, M. and Weeraratna, A. T. (2020). How the ageing microenvironment influences tumour progression.Nature Reviews Cancer, 20(2):89–106.

Firsanov, D., Zacher, M., Tian, X., Sformo, T. L., Zhao, Y., Tombline, G., Lu, J. Y., Zheng, Z., Perelli,L., Gurreri, E., Zhang, L., Guo, J., Korotkov, A., Volobaev, V., Biashad, S. A., Zhang, Z., Heid, J.,Maslov, A. Y., Sun, S., Wu, Z., Gigas, J., Hillpot, E. C., Martinez, J. C., Lee, M., Williams, A., Gilman, A., Hamilton, N., Strelkova, E., Haseljic, E., Patel, A., Straight, M. E., Miller, N., Ablaeva, J., Tam,L. M., Couderc, C., Hoopmann, M. R., Moritz, R. L., Fujii, S., Pelletier, A., Hayman, D. J., Liu, H., Cai, Y., Leung, A. K. L., Zhang, Z., Nelson, C. B., Abegglen, L. M., Schiffman, J. D., Gladyshev, V. N., Maley, C. C., Modesti, M., Genovese, G., Simons, M. J. P., Vijg, J., Seluanov, A., and Gorbunova, V. (2025). Evidence for improved DNA repair in long-lived bowhead whale. Nature.

Forsberg, L. A., Gisselsson, D., and Dumanski, J. P. (2017). Mosaicism in health and disease — clones picking up speed. Nature Reviews Genetics, 18(2):128–142.

Franceschi, C., Garagnani, P., Morsiani, C., Conte, M., Santoro, A., Grignolio, A., Monti, D., Capri, M., and Salvioli, S. (2018). The Continuum of Aging and Age-Related Diseases: Common Mechanisms but Different Rates. Frontiers in Medicine, 5:61.

Franco, I., Revêchon, G., and Eriksson, M. (2022). Challenges of proving a causal role of somatic mutations in the aging process. Aging Cell, 21(5):e13613.

Frank, S. A. (2005). Age-specific incidence of inherited versus sporadic cancers: A test of the multistage theory of carcinogenesis. Proceedings of the National Academy of Sciences, 102(4):1071–1075.

Frank, S. A. (2007). Dynamics of Cancer: Incidence, Inheritance, and Evolution. Princeton Series in Evolutionary Biology. Princeton University Press, Princeton, N.J.

Frank, S. A. (2010). Somatic evolutionary genomics: Mutations during development cause highly variable genetic mosaicism with risk of cancer and neurodegeneration. Proceedings of the National Academy of Sciences, 107(suppl_1):1725–1730.

Frank, S. A. (2014). Somatic Mosaicism and Disease. Current Biology, 24(12):R577–R581.

Frank, S. A. (2023). Robustness and complexity. Cell Systems, 14(12):1015–1020.

Frank, S. A. and Yanai, I. (2024). The origin of novel traits in cancer. Trends in Cancer, 10(10):880–892.

Freitas, A. A. and De Magalhães, J. P. (2011). A review and appraisal of the DNA damage theory of ageing. Mutation Research/Reviews in Mutation Research, 728(1-2)12–22.

Geritz, S. A. H., Kisdi, É., Meszéna, G., and Metz, J. A. J. (1998). Evolutionarily singular strategies and the adaptive growth and branching of the evolutionary tree. Evolutionary Ecology, 12(1):35–57.

Gerstung, M., Jolly, C., Leshchiner, I., Dentro, S. C., Gonzalez, S., Rosebrock, D., Mitchell, T. J., Rubanova, Y., Anur, P., Yu, K., Tarabichi, M., Deshwar, A., Wintersinger, J., Kleinheinz, K., Vázquez-García, I., Haase, K., Jerman, L., Sengupta, S., Macintyre, G., Malikic, S., Donmez, N., Livitz, D. G., Cmero, M., Demeulemeester, J., Schumacher, S., Fan, Y., Yao, X., Lee, J., Schlesner, M., Boutros, P. C., Bowtell, D. D., Zhu, H., Getz, G., Imielinski, M., Beroukhim, R., Sahinalp, S. C., Ji, Y., Peifer, M., Markowetz, F., Mustonen, V., Yuan, K., Wang, W., Morris, Q. D., Spellman, P. T., Wedge, D. C., Van Loo, P., PCAWG Evolution & Heterogeneity Working Group, and PCAWG Consortium (2020). The evolutionary history of 2,658 cancers. Nature, 578(7793):122–128.

Gladyshev, V. N. (2016). Aging: Progressive decline in fitness due to the rising deleteriome adjusted by genetic, environmental, and stochastic processes. Aging Cell, 15(4):594–602.

Goldsby, H. J., Knoester, D. B., Ofria, C., and Kerr, B. (2014). The Evolutionary Origin of Somatic Cells under the Dirty Work Hypothesis. PLoS Biology, 12(5):e1001858.

Gomes, N. M. V., Ryder, O. A., Houck, M. L., Charter, S. J., Walker, W., Forsyth, N. R., Austad, S. N., Venditti, C., Pagel, M., Shay, J. W., and Wright, W. E. (2011). Comparative biology of mammalian telomeres: Hypotheses on ancestral states and the roles of telomeres in longevity determination. Aging Cell, 10(5):761–768.

Gorbunova, V., Seluanov, A., Mao, Z., and Hine, C. (2007). Changes in DNA repair during aging. Nucleic Acids Research, 35(22):7466–7474.

Gorbunova, V., Seluanov, A., Zhang, Z., Gladyshev, V. N., and Vijg, J. (2014). Comparative genetics of longevity and cancer: Insights from long-lived rodents. Nature Reviews Genetics, 15(8):531–540.

Grant, P. R. and Grant, B. R. (1993). The Beak of the Finch: A Story of Evolution in Our Time. Princeton University Press, Princeton, NJ.

Guo, J., Huang, X., Dou, L., Yan, M., Shen, T., Tang, W., and Li, J. (2022). Aging and aging-related diseases: From molecular mechanisms to interventions and treatments. Signal Transduction and Targeted Therapy, 7(1):391.

Hall, J. (1988). Review and hypotheses: Somatic mosaicism: Observations related to clinical genetics. Am J Hum Genet., 43(4):355–363.

Hall, M. W. J., Jones, P. H., and Hall, B. A. (2019). Relating evolutionary selection and mutant clonal dynamics in normal epithelia. Journal of The Royal Society Interface, 16(156):20190230.

Hanahan, D. (2022). Hallmarks of Cancer: New Dimensions. Cancer Discovery, 12(1):31–46.

Harrington, J. S., Ryter, S. W., Plataki, M., Price, D. R., and Choi, A. M. K. (2023). Mitochondria in health, disease, and aging. Physiological Reviews, 103(4):2349–2422.

Healy, K., Guillerme, T., Finlay, S., Kane, A., Kelly, S. B. A., McClean, D., Kelly, D. J., Donohue, I., Jackson, A. L., and Cooper, N. (2014). Ecology and mode-of-life explain lifespan variation in birds and mammals. Proceedings of the Royal Society B: Biological Sciences, 281(1784):20140298.

Herms, A. and Jones, P. H. (2023). Somatic Mutations in Normal Tissues: New Perspectives on Early Carcinogenesis. Annual Review of Cancer Biology, 7(1):189–205.

Hochberg, M. E., Michalakis, Y., and De Meeus, T. (1992). Parasitism as a constraint on the rate of life-history evolution. Journal of Evolutionary Biology, 5(3):491–504.

Hochberg, M. E. and Noble, R. J. (2017). A framework for how environment contributes to cancer risk.Ecology Letters, 20(2):117–134.

Hochberg, M. E., Noble, R. J., and Braude, S. (2016). A Hypothesis to Explain Cancers in Confined Colonies of Naked Mole Rats. BioRxiv [Preprint].

Hochberg, M. E., Thomas, F., Assenat, E., and Hibner, U. (2013). Preventive Evolutionary Medicine of Cancers. Evolutionary Applications, 6(1):134–143.

Jaiswal, S. and Ebert, B. L. (2019). Clonal hematopoiesis in human aging and disease. Science, 366(6465):eaan4673.

Johnson, A. A., Shokhirev, M. N., and Shoshitaishvili, B. (2019). Revamping the evolutionary theories of aging. Ageing Research Reviews, 55:100947.

Kakiuchi, N. and Ogawa, S. (2021). Clonal expansion in non-cancer tissues. Nature Reviews Cancer, 21(4):239–256.

Kapsetaki, S. E., Compton, Z. T., Dolan, J., Harris, V. K., Mellon, W., Rupp, S. M., Duke, E. G., Harrison, T. M., Aksoy, S., Giraudeau, M., Vincze, O., McGraw, K. J., Aktipis, A., Tollis, M., Boddy, A. M., and Maley, C. C. (2024). Life history traits and cancer prevalence in birds. Evolution, Medicine, and Public Health, 12(1):105–116.

Kennedy, S. R., Loeb, L. A., and Herr, A. J. (2012). Somatic mutations in aging, cancer and neurodegeneration. Mechanisms of Ageing and Development, 133(4):118–126.

Kirkwood, T. and Holliday, R. (1979). The evolution of ageing and longevity. Proceedings of the Royal Society of London. Series B. Biological Sciences, 205(1161):531–546.

Kirkwood, T. B. and Rose, M. R. (1991). Evolution of senescence: Late survival sacrificed for reproduction. Philosophical Transactions of the Royal Society of London. Series B, Biological Sciences, 332(1262):15– 24.

Kirkwood, T. B. and Shanley, D. P. (2005). Food restriction, evolution and ageing. Mechanisms of Ageing and Development, 126(9):1011–1016.

Kirkwood, T. B. L. (2005). Understanding the Odd Science of Aging. Cell, 120(4):437–447.

Kokko, H. and Hochberg, M. E. (2015). Towards cancer-aware life-history modelling. Philosophical Transactions of the Royal Society B: Biological Sciences, 370(1673):20140234.

Kopinski, P. K., Singh, L. N., Zhang, S., Lott, M. T., and Wallace, D. C. (2021). Mitochondrial DNA variation and cancer. Nature Reviews Cancer, 21(7):431–445.

Krakauer, D. C. and Mira, A. (1999). Mitochondria and germ-cell death. Nature, 400(6740):125–126.

Kuijk, E., Blokzijl, F., Jager, M., Besselink, N., Boymans, S., Chuva De Sousa Lopes, S. M., Van Boxtel, R., and Cuppen, E. (2019). Early divergence of mutational processes in human fetal tissues. Science Advances, 5(5):eaaw1271.

Laconi, E., Marongiu, F., and DeGregori, J. (2020). Cancer as a disease of old age: Changing mutational and microenvironmental landscapes. British Journal of Cancer, 122(7):943–952.

Lakatos, E., Gunasri, V., Zapata, L., Househam, J., Heide, T., Trahearn, N., Swinyard, O., Cisneros, L., Lynn, C., Mossner, M., Kimberley, C., Spiteri, I., Cresswell, G. D., Llibre-Palomar, G., Mitchison,M., Maley, C. C., Jansen, M., Rodriguez-Justo, M., Bridgewater, J., Baker, A.-M., Sottoriva, A., and Graham, T. A. (2025). Epigenetically driven and early immune evasion in colorectal cancer evolution. Nature Genetics.

Lee-Six, H., Olafsson, S., Ellis, P., Osborne, R. J., Sanders, M. A., Moore, L., Georgakopoulos, N., Torrente, F., Noorani, A., Goddard, M., Robinson, P., Coorens, T. H. H., O’Neill, L., Alder, C., Wang, J., Fitzgerald, R. C., Zilbauer, M., Coleman, N., Saeb-Parsy, K., Martincorena, I., Campbell, P. J.,and Stratton, M. R. (2019). The landscape of somatic mutation in normal colorectal epithelial cells.Nature, 574(7779):532–537.

Li, Q., Wang, S., Milot, E., Bergeron, P., Ferrucci, L., Fried, L. P., and Cohen, A. A. (2015). Homeostatic dysregulation proceeds in parallel in multiple physiological systems. Aging Cell, 14(6):1103–1112.

Lynch, M. (2010a). Evolution of the mutation rate. Trends in Genetics, 26(8):345–352.

Lynch, M. (2010b). Rate, molecular spectrum, and consequences of human mutation. Proceedings of the National Academy of Sciences, 107(3):961–968.

Lynch, M., Ackerman, M. S., Gout, J.-F., Long, H., Sung, W., Thomas, W. K., and Foster, P. L. (2016). Genetic drift, selection and the evolution of the mutation rate. Nature Reviews Genetics, 17(11):704–714.

MacRae, S. L., Zhang, Q., Lemetre, C., Seim, I., Calder, R. B., Hoeijmakers, J., Suh, Y., Gladyshev,V. N., Seluanov, A., Gorbunova, V., Vijg, J., and Zhang, Z. D. (2015). Comparative analysis of genome maintenance genes in naked mole rat, mouse, and human. Aging Cell, 14(2):288–291.

Maklakov, A. A. and Chapman, T. (2019). Evolution of ageing as a tangle of trade-offs: Energy versus function. Proceedings of the Royal Society B: Biological Sciences, 286(1911):20191604.

Manders, F., Van Boxtel, R., and Middelkamp, S. (2021). The Dynamics of Somatic Mutagenesis During Life in Humans. Frontiers in Aging, 2:802407.

Marongiu, F. and DeGregori, J. (2022). The sculpting of somatic mutational landscapes by evolutionary forces and their impacts on aging-related disease. Molecular Oncology, 16(18):3238–3258.

Martin, P., Pardo-Pastor, C., Jenkins, R. G., and Rosenblatt, J. (2024). Imperfect wound healing sets the stage for chronic diseases. Science, 386(6726):eadp2974.

Martincorena, I. (2019). Somatic mutation and clonal expansions in human tissues. Genome Medicine, 11(1):35.

Martincorena, I. and Campbell, P. J. (2015). Somatic mutation in cancer and normal cells. Science, 349(6255):1483–1489.

Martincorena, I., Roshan, A., Gerstung, M., Ellis, P., Van Loo, P., McLaren, S., Wedge, D. C., Fullam, A., Alexandrov, L. B., Tubio, J. M., Stebbings, L., Menzies, A., Widaa, S., Stratton, M. R., Jones,P. H., and Campbell, P. J. (2015). High burden and pervasive positive selection of somatic mutations in normal human skin. Science, 348(6237):880–886.

Medawar, P. (1952). An Unsolved Problem of Biology: An Inaugural Lecture Delivered at University College, London, 6 December, 1951. H.K. Lewis and Company.

Medzhitov, R., Schneider, D. S., and Soares, M. P. (2012). Disease Tolerance as a Defense Strategy.Science, 335(6071):936–941.

Milholland, B., Suh, Y., and Vijg, J. (2017). Mutation and catastrophe in the aging genome. Experimental Gerontology, 94:34–40.

Moore, L., Cagan, A., Coorens, T. H. H., Neville, M. D. C., Sanghvi, R., Sanders, M. A., Oliver, T.R. W., Leongamornlert, D., Ellis, P., Noorani, A., Mitchell, T. J., Butler, T. M., Hooks, Y., Warren,A. Y., Jorgensen, M., Dawson, K. J., Menzies, A., O’Neill, L., Latimer, C., Teng, M., Van Boxtel, R., Iacobuzio-Donahue, C. A., Martincorena, I., Heer, R., Campbell, P. J., Fitzgerald, R. C., Stratton,M. R., and Rahbari, R. (2021). The mutational landscape of human somatic and germline cells. Nature, 597(7876):381–386.

Mustjoki, S. and Young, N. S. (2021). Somatic Mutations in “Benign” Disease. New England Journal of Medicine, 384(21):2039–2052.

Niccoli, T. and Partridge, L. (2012). Ageing as a Risk Factor for Disease. Current Biology, 22(17):R741– R752.

Niedernhofer, L. J., Gurkar, A. U., Wang, Y., Vijg, J., Hoeijmakers, J. H., and Robbins, P. D. (2018). Nuclear Genomic Instability and Aging. Annual Review of Biochemistry, 87(1):295–322.

Noble, R., Kaltz, O., and Hochberg, M. E. (2015). Peto’s paradox and human cancers. Philosophical Transactions of the Royal Society B: Biological Sciences, 370(1673):20150104.

Nowak, M. A., Michor, F., and Iwasa, Y. (2003). The linear process of somatic evolution. Proceedings of the National Academy of Sciences of the United States of America, 100(25):14966–14969.

Nunney, L. (2003). The population genetics of multistage carcinogenesis. Proceedings of the Royal Society of London. Series B: Biological Sciences, 270(1520):1183–1191.

Nunney, L. (2013). The real war on cancer: The evolutionary dynamics of cancer suppression. Evolutionary Applications, 6(1):11–19.

Olafsson, S. and Anderson, C. A. (2021). Somatic mutations provide important and unique insights into the biology of complex diseases. Trends in Genetics, 37(10):872–881.

Omholt, S. W. and Kirkwood, T. B. L. (2021). Aging as a consequence of selection to reduce the environmental risk of dying. Proceedings of the National Academy of Sciences, 118(22):e2102088118.

Osorio, F. G., Rosendahl Huber, A., Oka, R., Verheul, M., Patel, S. H., Hasaart, K., De La Fonteijne, L., Varela, I., Camargo, F. D., and Van Boxtel, R. (2018). Somatic Mutations Reveal Lineage Relationships and Age-Related Mutagenesis in Human Hematopoiesis. Cell Reports, 25(9):2308–2316.e4.

Pepke, M. L. and Eisenberg, D. T. A. (2022). On the comparative biology of mammalian telomeres: Telomere length co-evolves with body mass, lifespan and cancer risk. Molecular Ecology, 31(23):6286–6296.

Pomatto, L. C. D. and Davies, K. J. A. (2017). The role of declining adaptive homeostasis in ageing.The Journal of Physiology, 595(24):7275–7309.

Promislow, D., Anderson, R. M., Scheffer, M., Crespi, B., DeGregori, J., Harris, K., Horowitz, B. N., Levine, M. E., Riolo, M. A., Schneider, D. S., Spencer, S. L., Valenzano, D. R., and Hochberg, M. E. (2022). Resilience integrates concepts in aging research. iScience, 25(5):104199.

Promislow, D. E. (1994). DNA Repair and the Evolution of Longevity: A Critical Analysis. Journal of Theoretical Biology, 170(3):291–300.

Proukakis, C. (2020). Somatic mutations in neurodegeneration: An update. Neurobiology of Disease, 144:105021.

Risques, R. A. and Promislow, D. E. L. (2018). All’s well that ends well: Why large species have short telomeres. Philosophical Transactions of the Royal Society B: Biological Sciences, 373(1741):20160448.

Rockweiler, N. B., Ramu, A., Nagirnaja, L., Wong, W. H., Noordam, M. J., Drubin, C. W., Huang, N.,Miller, B., Todres, E. Z., Vigh-Conrad, K. A., Zito, A., Small, K. S., Ardlie, K. G., Cohen, B. A., and Conrad, D. F. (2023). The origins and functional effects of postzygotic mutations throughout the human life span. Science, 380(6641):eabn7113.

Rozhok, A. and DeGregori, J. (2019). Somatic maintenance impacts the evolution of mutation rate. BMC Evolutionary Biology, 19(1):172.

Rozhok, A. I. and DeGregori, J. (2016). The Evolution of Lifespan and Age-Dependent Cancer Risk.Trends in Cancer, 2(10):552–560.

Rozhok, A. I. and DeGregori, J. (2020). The three dimensions of somatic evolution: Integrating the role of genetic damage, life-history traits, and aging in carcinogenesis. Evolutionary Applications, 13(7):1569–1580.

Schumacher, B., Pothof, J., Vijg, J., and Hoeijmakers, J. H. J. (2021). The central role of DNA damage in the ageing process. Nature, 592(7856):695–703.

Seluanov, A., Gladyshev, V. N., Vijg, J., and Gorbunova, V. (2018). Mechanisms of cancer resistance in long-lived mammals. Nature Reviews Cancer, 18(7):433–441.

Shen, X., Wang, C., Zhou, X., Zhou, W., Hornburg, D., Wu, S., and Snyder, M. P. (2024). Nonlinear dynamics of multiomics profiles during human aging. Nature Aging, 4:1619–1634.

St Sauver, J. L., Boyd, C. M., Grossardt, B. R., Bobo, W. V., Finney Rutten, L. J., Roger, V. L., Ebbert,J. O., Therneau, T. M., Yawn, B. P., and Rocca, W. A. (2015). Risk of developing multimorbidity across all ages in an historical cohort study: Differences by sex and ethnicity. BMJ Open, 5(2):e006413.

Stearns, S. C. (1987). The selection-arena hypothesis. In Stearns, S. C., editor, The Evolution of Sex and Its Consequences, volume 55, pages 337–349. Birkhäuser Basel, Basel.

Strogatz, S. H. (2018). Nonlinear Dynamics and Chaos: With Applications to Physics, Biology, Chemistry, and Engineering. Westview Press, 2nd edition.

Szathmáry, E. (2015). Toward major evolutionary transitions theory 2.0. Proceedings of the National Academy of Sciences of the United States of America, 112(33):10104–10111.

Tian, X., Doerig, K., Park, R., Can Ran Qin, A., Hwang, C., Neary, A., Gilbert, M., Seluanov, A., and Gorbunova, V. (2018). Evolution of telomere maintenance and tumour suppressor mechanisms across mammals. Philosophical Transactions of the Royal Society B: Biological Sciences, 373(1741):20160443.

Tian, X., Seluanov, A., and Gorbunova, V. (2017). Molecular Mechanisms Determining Lifespan in Short- and Long-Lived Species. Trends in Endocrinology & Metabolism, 28(10):722–734.

Tollis, M., Schneider-Utaka, A. K., and Maley, C. C. (2020). The evolution of human cancer gene duplications across mammals. Molecular Biology and Evolution, 37(10):2875–2886.

Vijg, J. (2021). From DNA damage to mutations: All roads lead to aging. Ageing Research Reviews, 68:101316.

Vijg, J. and Dong, X. (2020). Pathogenic Mechanisms of Somatic Mutation and Genome Mosaicism in Aging. Cell, 182(1):12–23.

Vincze, O., Colchero, F., Lemaître, J.-F., Conde, D. A., Pavard, S., Bieuville, M., Urrutia, A. O., Ujvari, B., Boddy, A. M., Maley, C. C., Thomas, F., and Giraudeau, M. (2022). Cancer risk across mammals. Nature, 601(7892):263–267.

Weeden, C. E., Hill, W., Lim, E. L., Grönroos, E., and Swanton, C. (2023). Impact of risk factors on early cancer evolution. Cell, 186(8):1541–1563.

Williams, G. C. (1957). Pleiotropy, Natural Selection, and the Evolution of Senescence. Evolution, 11(4):398–411.

Wright, A., Charlesworth, B., Rudan, I., Carothers, A., and Campbell, H. (2003). A polygenic basis for late-onset disease. Trends in Genetics, 19(2):97–106.

Wu, S., Zhu, W., Thompson, P., and Hannun, Y. A. (2018). Evaluating intrinsic and non-intrinsic cancer risk factors. Nature Communications, 9(1):3490.

Youssoufian, H. and Pyeritz, R. E. (2002). Mechanisms and consequences of somatic mosaicism in humans.Nature Reviews Genetics, 3(10):748–758.

Zhang, L., Dong, X., Tian, X., Lee, M., Ablaeva, J., Firsanov, D., Lee, S.-G., Maslov, A. Y., Gladyshev,V. N., Seluanov, A., Gorbunova, V., and Vijg, J. (2021). Maintenance of genome sequence integrity in long- and short-lived rodent species. Science Advances, 7(44):eabj3284.

Zhu, M., Lu, T., Jia, Y., Luo, X., Gopal, P., Li, L., Odewole, M., Renteria, V., Singal, A. G., Jang, Y.,Ge, K., Wang, S. C., Sorouri, M., Parekh, J. R., MacConmara, M. P., Yopp, A. C., Wang, T., and Zhu, H. (2019). Somatic Mutations Increase Hepatic Clonal Fitness and Regeneration in Chronic Liver Disease. Cell, 177(3):608–621.e12.

