## Supplementary Information for "Disease as a Mediator of Somatic Mutation - Lifespan Coevolution: Model Analysis and Empirical Tests"

### Supplementary Information. Local Stability and Dynamics

We analyze the local stability and local return dynamics of the system

$$\dot{M} = aQ - dMF, \qquad \dot{F} = vQ - gF - rMF. \tag{S1}$$

Here $M$ denotes somatic mutation rate (SMR), $F$ denotes lifespan, and the parameters satisfy

$$Q > 0, \qquad a > 0, \qquad d > 0, \qquad v > 0, \qquad g > 0, \tag{S2}$$

while $r$ may be positive or negative after the parameter convention $r - l \to r$ (see main text). In what follows, $M^*$ and $F^*$ denote the positive interior equilibrium, and $S = dv - ar$.

**Local Stability**. The local stability analysis requires the Jacobian matrix, $J^*$,

$$J^* = \begin{pmatrix} -dF^* & -dM^* \\ -rF^* & -(rM^* + g) \end{pmatrix}, \tag{S3}$$

together with its trace $T$, determinant $D$, and discriminant $\Delta$:

$$T = \mathrm{tr}(J^*) = -\frac{QS}{g} - g - \frac{agr}{S}, \qquad D = \det(J^*) = QS, \qquad \Delta = T^2 - 4D. \tag{S4}$$

Whenever $S > 0$, we have $D > 0$ and $T < 0$. Therefore the interior equilibrium is always locally asymptotically stable whenever it exists. Small perturbations away from equilibrium therefore decay back toward the equilibrium, either through damped oscillatory transients or through a non-oscillatory monotone return.

**Parameter Conditions for Damped Oscillations**. For fixed $Q$, $a$, $d$, $v$, and $g$, and with $S$ varied through $r$, the boundary between non-oscillatory and oscillatory return is defined by

$$\frac{QS}{g} + \frac{gdv}{S} = 2\sqrt{QS}. \tag{S5}$$

Let

$$x = \sqrt{QS}. \tag{S6}$$

The boundary condition becomes

$$\frac{x^2}{g} + \frac{gdvQ}{x^2} = 2x, \tag{S7}$$

or equivalently

$$x^4 - 2g\,x^3 + g^2 dvQ = 0. \tag{S8}$$

This quartic has two distinct positive roots $x_- < x_+$ iff

$$g^2 > \frac{16}{27} Qdv. \tag{S9}$$

If (S9) holds, then an oscillatory interval exists. If this inequality is not satisfied, then all local return trajectories are non-oscillatory. A simpler sufficient, but not necessary, condition for the existence of such an oscillatory interval is

$$g > \sqrt{Qdv}. \tag{S10}$$

Therefore, such an interval is promoted (but not ensured) for short lived species with high somatic mutation rates.

**Return Time to Equilibrium**. The characteristic polynomial from the Jacobian is

$$\lambda^2 - T\lambda + D = 0, \tag{S11}$$

and hence the eigenvalues are

$$\lambda_{1,2} = \frac{T \pm \sqrt{\Delta}}{2}. \tag{S12}$$

The eigenvalues have negative real parts whenever the interior equilibrium exists. The asymptotic time for a generic small perturbation to shrink by a factor $e$ in the linearized dynamics is the *e-fold time* $\tau_e$, defined as

$$\tau_e := \frac{1}{|\Re(\lambda_{\text{slow}})|}, \tag{S13}$$

where $\lambda_{\text{slow}}$ is the eigenvalue whose real part is closest to 0. The e-fold time $\tau_e$ is the inverse of the slowest exponential decay rate and is the characteristic time over which a perturbation decays to about 37% of its initial size (see main text). Biologically, larger $\tau_e$ means a longer transients after (implicit) environmental, demographic, or evolutionary perturbation. Smaller $\tau_e$ means faster restoration of the local SMR–lifespan steady state.

The expression for $\tau_e$ differs between oscillatory and non-oscillatory departures from equilibrium. It is useful to aggregate parameters, such that

$$B = -T = \frac{QS}{g} + \frac{gdv}{S}, \qquad D = QS. \tag{S14}$$

For a stable focus, corresponding to an oscillatory local return,

$$\tau_{\text{focus}} = \frac{2}{B} = \frac{2}{\frac{QS}{g} + \frac{gdv}{S}}. \tag{S15}$$

Smaller $Q$ increases $\tau_{\text{focus}}$, so perturbations decay more slowly in smaller-body-size systems. Increasing either $d$ or $v$ decreases $\tau_{\text{focus}}$. Thus, both stronger SMR suppression and stronger lifespan support speed recovery.

In contrast, the effects of $a$, $r$, and $g$ on $\tau_{\text{focus}}$ are context-dependent. The switch is determined by

$$K = QS^2 - dg^2v. \tag{S16}$$

When $K > 0$, the contribution $QS/g$ dominates the damping denominator, whereas, when $K < 0$, the contribution $gdv/S$ dominates. Thus, stronger mutation input ($a$) changes in net coupling $r$, and stronger lifespan loss ($g$) can either lengthen or shorten recovery time depending on which contribution dominates.

For a stable node, corresponding to non-oscillatory local return,

$$\tau_{\text{node}} = \frac{2}{B - \sqrt{B^2 - 4D}}. \tag{S17}$$

The stable node corresponds to non-oscillatory local recovery. Real eigenvalues do not guarantee that each state variable is monotone for every initial perturbation, so overshoot of an individual coordinate is not excluded. In this regime, the effects of parameters on return time are generally more context-dependent than in the focus case. Except for special cases, there are no general mappings of parameter values to return times as measured by $\tau_{\text{node}}$.
